# Archival influenza virus genomes from Europe reveal genomic and phenotypic variability during the 1918 pandemic

**DOI:** 10.1101/2021.05.14.444134

**Authors:** Livia Victoria Patrono, Bram Vrancken, Matthias Budt, Ariane Düx, Sebastian Lequime, Sengül Boral, M. Thomas P. Gilbert, Jan F. Gogarten, Luisa Hoffmann, David Horst, Kevin Merkel, David Morens, Baptiste Prepoint, Jasmin Schlotterbeck, Verena Schuenemann, Marc A. Suchard, Jeffery K. Taubenberger, Luisa Tenkhoff, Christian Urban, Navena Widulin, Eduard Winter, Michael Worobey, Fabian H. Leendertz, Thomas Schnalke, Thorsten Wolff, Philippe Lemey, Sébastien Calvignac-Spencer

## Abstract

The 1918 influenza pandemic was the deadliest respiratory pandemic of the 20th century and determined the genomic make-up of subsequent human influenza A viruses (IAV). Here, we analyze the first 1918 IAV genomes from Europe and from the first, milder wave of the pandemic. 1918 IAV genomic diversity is consistent with local transmission and frequent long-distance dispersal events and *in vitro* polymerase characterization suggests potential phenotypic variability. Comparison of first and second wave genomes shows variation at two sites in the nucleoprotein gene associated with resistance to host antiviral response, pointing at a possible adaptation of 1918 IAV to humans. Finally, phylogenetic estimates based on extended molecular clock modelling suggests a pure pandemic descent of seasonal H1N1 IAV as an alternative to the hypothesis of an intrasubtype reassortment origin.

**One Sentence Summary:** Much can be learned about past pandemics by uncovering their footprints in medical archives, which we here demonstrate for the 1918 flu pandemic.

## Main Text

As the COVID-19 pandemic unfolds (*1*), we are facing many questions regarding the emergence and spread of a novel pathogen in human populations. Although every pandemic is unique and making meaningful comparisons is challenging, the current situation naturally rekindles interest in past pandemics (*2*).

The 1918 influenza A (H1N1) pandemic (hereafter 1918 pandemic) was the largest global catastrophe of infectious origin to affect humankind in the last century. The disease proceeded in three waves: a mild one in the spring of 1918, followed by deadlier waves in the 1918 fall and 1919 winter (*3*). Beyond mortality records from that period that allow for comparisons of the scale of this pandemic with subsequent ones, very little is known about even basic biological consequences of the fast and global spread of the 1918 virus. For example, it remains essentially unknown how much genomic diversity arose during the 1918 pandemic and how this diversity impacted viral phenotypes.

In the late 1990s, molecular analyses of formalin-fixed, paraffin-embedded tissue blocks and permafrost-preserved bodies from victims of the 1918 pandemic (*4*) ascertained that the causative agent was an influenza A virus (IAV) of the H1N1 subtype (*5*). The reconstruction of two complete IAV genomes from Brevig Mission, Alaska (hereafter BM) (*6*) and Camp Upton, New York (hereafter CU) (*7*), as well as the identification of a few other partial sequences (*8, 9*) informed evolutionary and functional investigations. These revealed that the 1918 virus harbored gene segments drawn from the diversity of IAV strains circulating in an avian reservoir (*10*), and that the hemagglutinin (HA) and polymerase complex genes were major determinants of its pathogenicity (*11, 12*). However, our knowledge about an agent that caused an estimated 50-100 million deaths (*13*) remains extremely limited, notably because it relies on a remarkably sparse corpus of genomic/genetic data, the majority stemming from North America.

We have started to meet this gap by applying recent advances in nucleic acid recovery from archival samples (*7, 14*) to as-yet unexplored European pathology collection material, including the medical archive started by Rudolf Virchow in mid 1800s Germany (*15*). This enabled us to sequence complete and partial 1918 influenza virus genomes from specimens sampled in two German cities, then use these data to provide new insights into the genomic and phenotypic diversity of the pandemic strains.

### 1918 influenza virus genomes from Munich and Berlin

To explore the emergence, pandemic and post-pandemic periods, we selected 13 formalin-fixed lung specimens dated between 1900 and 1931, preserved within the Berlin Museum of Medical History at the Charité (Berlin, Germany, n=11) and the pathology collection (Narrenturm) of the Natural History Museum in Vienna, Austria (n=2). This set included six specimens collected during pandemic years in Europe (n=4 in 1918, n=2 in 1919). Details about all specimens, including initial diagnosis, are available in the supplementary information. For each specimen, we heat treated 200 mg of formalin-fixed lung tissue to reverse macromolecule cross-links induced by formalin, then performed nucleic acid extraction. Following DNase treatment and ribosomal RNA depletion, we built high-throughput sequencing libraries and shotgun sequenced them on Illumina® platforms.

We identified IAV reads in libraries from 3 of 13 specimens, all of which date to 1918 and were associated with histopathological findings of bronchopneumonia (**Fig. 1A**; **supplementary text S1 and Fig. S1**). Viral RNA preservation was generally good, as determined by median fragment lengths well above 100 nucleotides and the lack of any obvious damage pattern (**Fig. S9-S11**). Human RNA fragments were on average shorter, possibly indicating better protection of encapsidated viral RNA (**supplementary text S2**). We did not detect any other viral agent likely to be involved in severe respiratory disease, but identified a number of bacteria which have been previously associated with respiratory diseases (e.g. *Pseudomonas aeruginosa* and *Mycobacterium kansasii* in specimens that did not produce influenza reads) and with 1918 influenza (*Staphylococcus aureus, Streptococcus pneumoniae, Klebsiella pneumoniae, Pasteurella multocida*); in 2 specimens bacterial colonization was also evident from histopathology **(supplementary text S1 and Fig. S1)**.

**Fig. 1:**
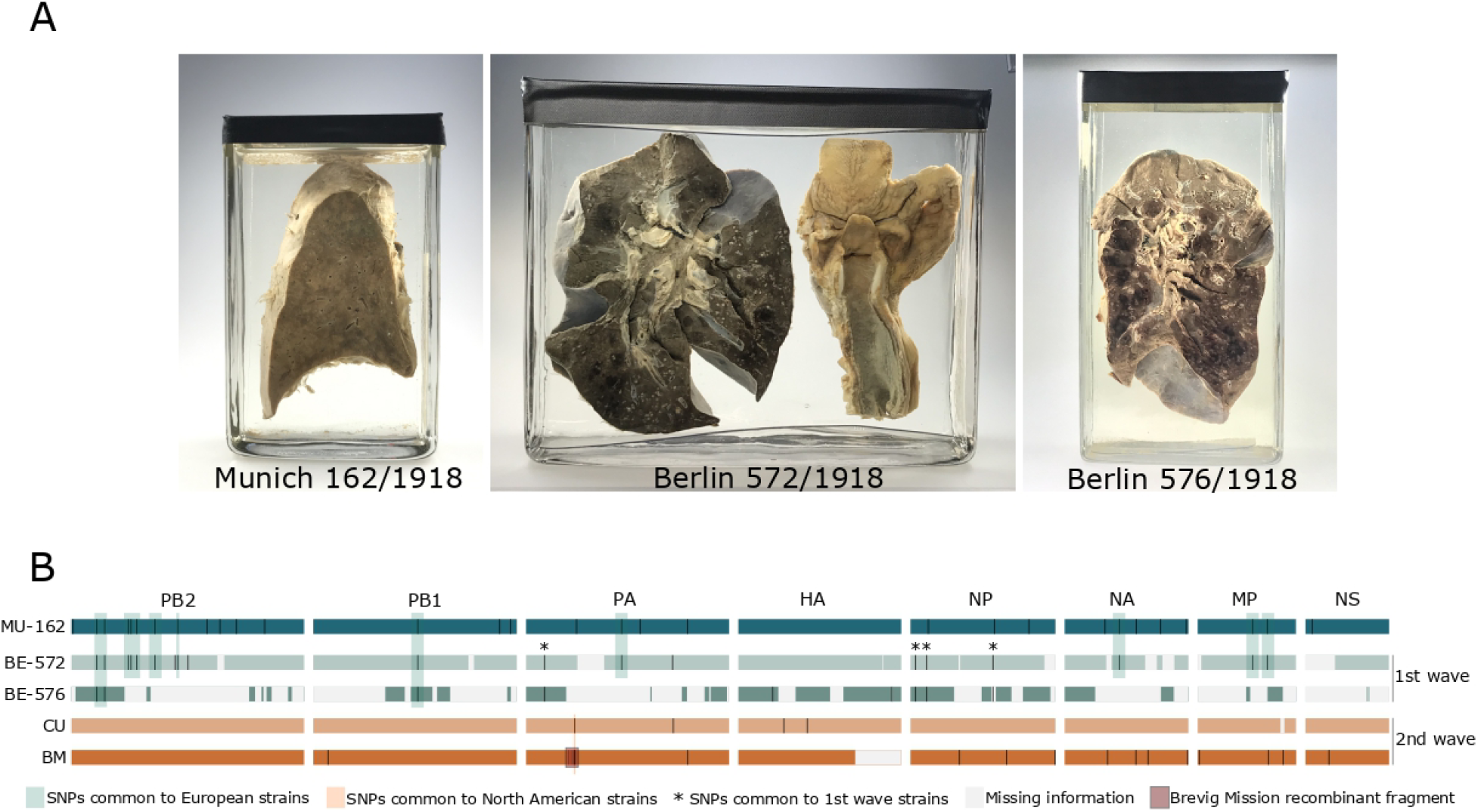
Specimens positive for influenza A virus (A) and comparison of all available 1918 influenza A virus genomes (B). We identified single nucleotide polymorphisms (SNPs) in the new genomes using BM as a reference; for BM and CU, we plotted SNPs unique to these genomes. Missing information represents areas where we did not get any coverage or this was lower than our consensus calling criteria.

Increased sequencing efforts for the three influenza-positive samples MU-162 (Munich), BE-572 and BE-576 (Berlin) yielded 40.133.161, 31.989.479 and 14.965.377 high quality reads, respectively. Of these, 55.5%, 0.33% and 0.1% mapped to the influenza virus strain A/Brevig Mission/1/1918, which allowed us to reconstruct a complete 1918 influenza virus genome at a 1944x average coverage depth for MU-162, as well as significant portions of the genomes for BE-572 and BE-576 (89.3% and 57.2%, respectively).

### Genomic comparison and phylodynamics of 1918 influenza viruses

Together with the available BM and CU sequences, we used these influenza genomes from Germany to assess the genomic diversity of: (i) strains simultaneously involved in local transmission (BE-572 and BE-576), (ii) strains circulating in Europe (MU-162, BE-572 and BE- 576) and North America (BM and CU), (iii) strains circulating during the first (BE-572 and BE- 576) and second wave (BM and CU) of the pandemic.

The two Berlin genomes sampled on June 28^th^ 1918, for which only a portion of the genome could be compared, differed from each other at only two (non-synonymous) nucleotide positions in the HA gene (**Fig. 1B**). These positions were however polymorphic in BE-576, with the minor variant being identical to BE-572. At a country/continent scale, BE-572 and MU-162 on one hand and BM and CU on the other differed both by 22 and 15 single nucleotide polymorphisms (SNPs; 11 non-synonymous, all genes but HA, NS and MP and 7 non-synonymous, all genes but NP and PB2, respectively, **Table 1 and S4**; **Fig. 1B**). The four pairwise comparisons of European and North American strains identified 22-43 SNPs, with the two best genomes (MU-162 and BM) differing at 43 positions, 16 of which coding for amino acid changes (all genes but HA). When comparing the first wave European strain BE-572 with second wave strains BM and CU, we identified 29 and 22 SNPs (11 non-synonymous, all genes but HA and NS genes, and 8 non-synonymous, all genes but NS genes, respectively).

**Table 1.**
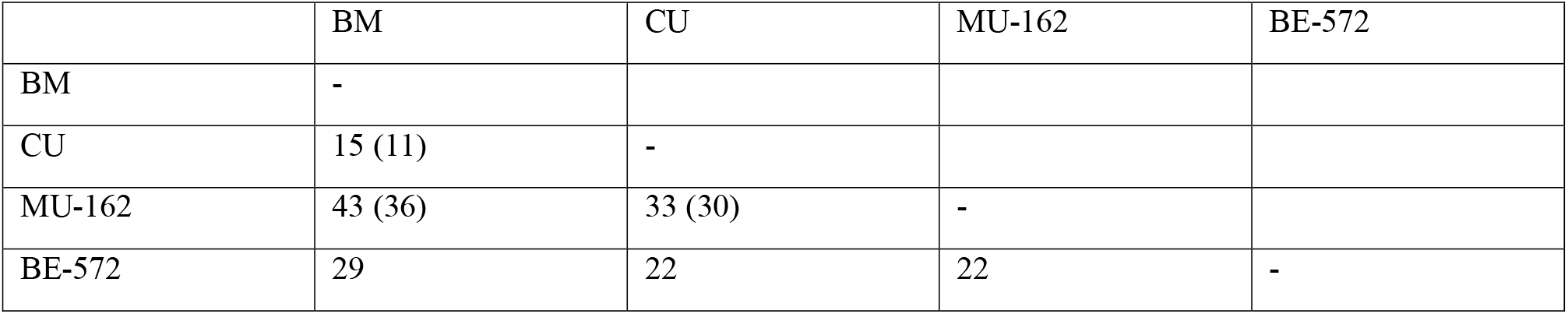
Number of differences between the highest quality genomes. Numbers in () represent the differences counted in the positions covered by BE-572 (12023 nt). BE-576 was not included in these comparisons because of its lower quality. A 69 nt fragment in PA with a signal of recombination (8 SNPs, 1 coding) was excluded from this comparison.

We could not fully disentangle these comparisons (the two Berlin genomes are not complete, and the two accurately dated genomes from the first wave are from European specimens, while the two accurately dated genomes from the second wave are from North America) and our sample size was very small. However, keeping these limitations in mind, it appears that within-outbreak variability in Berlin was negligible, while genomes sampled on the same continents (0.11-0.16%) and during the same wave (0.11%) exhibited lower overall divergence than genomes sampled on different continents (0.16-0.32%) and during different waves (0.16-0.21%). This pattern is compatible with spatio-temporal effects of local transmission.

To further investigate the question of 1918 virus spread, we took advantage of the availability of a total of 21 HA sequences sampled in Europe and North America during all pandemic waves to infer a time-scaled evolutionary history in a Bayesian phylogenetic framework (*16*) This reconstruction demonstrated an interspersed clustering of the three German sequences with North American strains, suggesting no geographic segregation between continents (**Fig. 2**). Similarly, two sequences sampled three months apart in London during the 1918-1919 winter waves of the pandemic (November 1918 and February 1919, respectively) clustered separately, albeit with low support (posterior probability .75 and .73, **Fig. 2 and S7**). Models in which monophyletic clustering was enforced on the European lineages showed a decisively lower model fit (ln Bayes factor [BF] ≥ 8.7, **supplementary text S6**), supporting the hypothesis of extensive, and most likely bi-directional, transatlantic movement of pandemic strains, in line with the historical context of human movement near the end of World War I. Altogether, genomic data support a scenario of predominantly local transmission with frequent long-distance dispersal events.

**Fig. 2:**
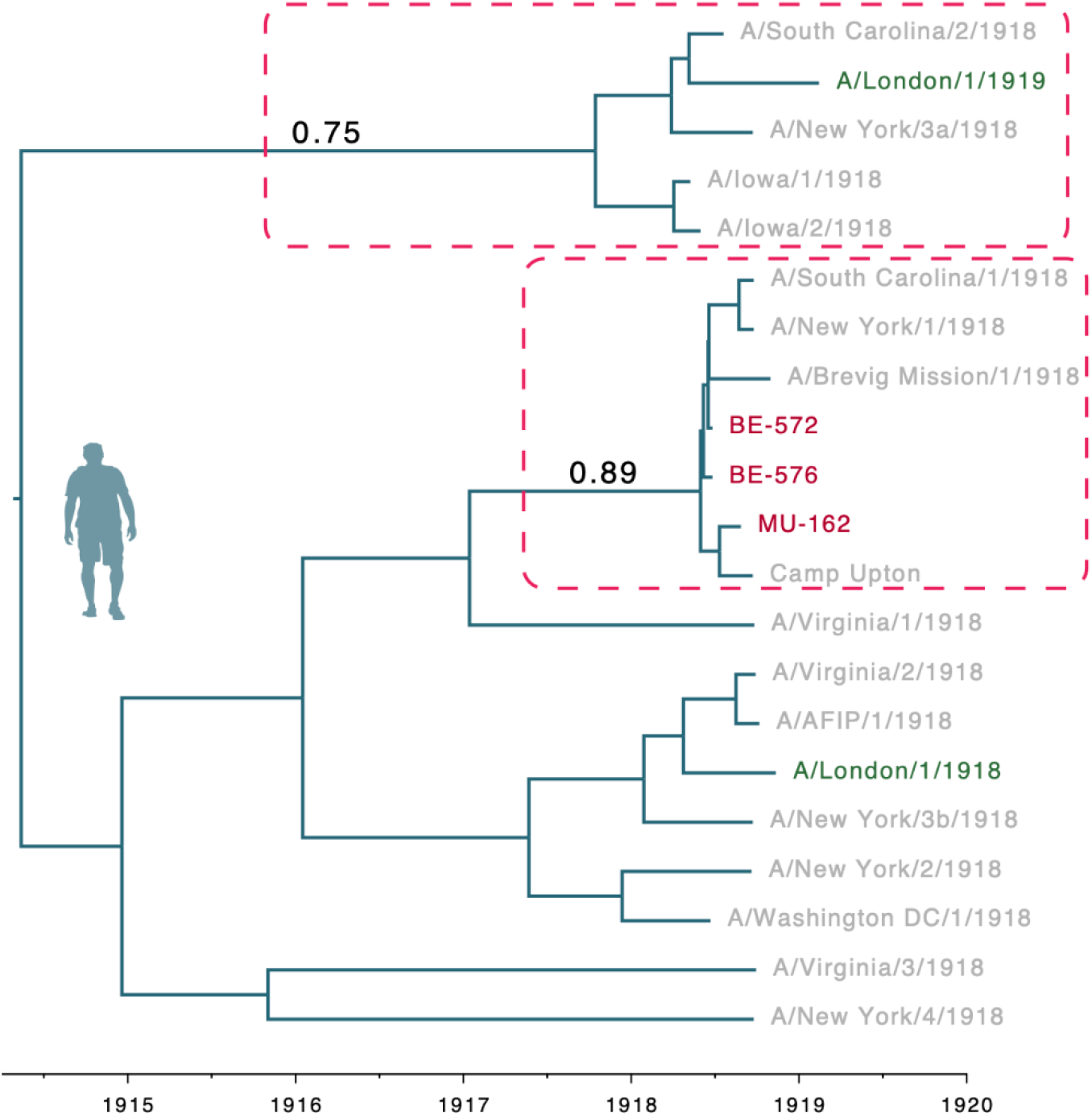
Maximum clade credibility time tree estimated for HA sequences from 1918 flu strains. US strains are in light grey pink, European strains are in dark red (Germany) and dark green (UK). Dashed rectangles highlight clades comprising strains from different continents; for these clades posterior probabilities are reported above stem branches. The BE-576 consensus genome with ambiguities was used in this analysis; an equivalent reconstruction using majority rule consensus base calling can be found in Fig. S7.

Finally, we identified a 69 nt stretch of the PA of BM comprising 8 SNPs that shows a high degree of identity to IAV strains circulating in 1933 or later (**Table S4**). Given the size of this fragment and that of the fragments initially targeted to characterize the BM PA sequence (*17*), this apparent recombination may reflect the contamination of one of the PCR reactions that allowed for the reconstruction of the BM genome with a PCR product derived from a later influenza A virus strain. However, we cannot formally exclude that the BM PA fragment is a genuine but rare case of natural recombination (*18*) or that it represents a sequencing artefact due to intrahost variation. Resequencing of the BM genome using PCR-independent methods should clarify this issue.

### Potential signatures of avian-to-human adaptation in 1918 influenza viruses

In light of the hypothesis of an avian origin of the pandemic virus, we investigated the presence of amino acid (aa) signatures known or suspected to be associated with avian-to-human adaptation, either because they have been characterized functionally or found to be distributed differentially across bird- and human-infecting influenza viruses. We first examined the high quality (but imprecisely dated) genome derived from MU-162 and identified two such positions. In PB2, we found a M631L aa change within the PB2 627 host range domain. Although this mutation has recently been described as a main mediator of adaptation and lethality of an avian influenza virus in mice (*19*), all but one human H1N1 strains (including all other 1918 influenza viruses) present a methionine at this position, suggesting that this change did not have profound evolutionary implications. On the contrary, we found that MU-162 has a leucine (L) at position 61 in NP, which is the residue most commonly found in human H1N1 strains prior to the 2009pdm (**Fig. S12**), whereas an overwhelming majority of avian influenza A viruses present an isoleucine (I) at this position (*20*). Interestingly, MU-162 was the only 1918 influenza virus presenting the I61L change, indicating this mutation only reached fixation after the first two waves of the pandemic.

We then considered potential coding variation between first and second wave strains. The second wave of the 1918 pandemic was indeed much more severe than the first, and potential adaptations of the virus have been proposed as one possible explanation, among others (*3*). The presence of the aa residue G222 in the receptor binding domain of the H1 subtype HA protein has already been discussed in this context (*8*). This residue confers a binding affinity for both avian and human glycans, while the human-like D222 only efficiently binds human glycans (*21, 22*). In a recent study, three of four first wave strain sequences presented G222 while only two of nine second wave strains did (all other strains presented D222) (*8*). The three German HAs sequenced here carried the human-like D222, which reduces the likelihood that the frequency of the two variants varied significantly between the first two pandemic waves.

The availability of genome-wide information about two first wave strains also allowed us to identify two other aa changes that potentially mark a difference between first and second wave strains. We detected these variations at sites of the NP known to influence host range and susceptibility/resistance to the interferon-induced MxA antiviral protein (*20, 23, 24*): first wave strains BE-572 and BE-576 carried avian-like residues at position 16 (G) and 283 (L), whereas the fall strains and MU-162 carried D16 and P283 found in most human H1N1 strains with the exception of the 2009 pandemic H1N1 virus (see references (*23–26*) and **Fig. S12**). Position 283 is located in the body of the NP and presence of a proline at this site confers resistance to the human MxA protein, to which D16 also contributes, albeit to a lower extent (*24, 26*). In addition to supporting the hypothesis of an avian origin of this gene, these mutations may represent hallmarks of early adaptation to humans: during the first months of the pandemic, 1918 influenza viruses may have evolved a better capacity to evade the innate interferon response, which is an important aspect of influenza virus pathogenicity.

### Comparison of *in vitro* polymerase activity of 1918 influenza viruses

To start exploring the phenotypic impact of the genomic differences observed *in vitro*, we focused on the MU-162 strain for which we obtained a high-quality complete genome sequence. Since the polymerase complex contained most of the coding mutations found in MU-162 (4 in PB2, 2 in PB1, 3 in PA and 1 in NP proteins), and is a major pathogenicity factor for BM, we resynthesized viral genes and performed a functional comparison of the activity of the BM and MU-162 polymerases after reconstitution in transfected cells. Using reporter assays, dose response curves revealed a two-fold higher activity of BM compared to MU-162 polymerase (**Fig. 3A**).

**Fig. 3:**
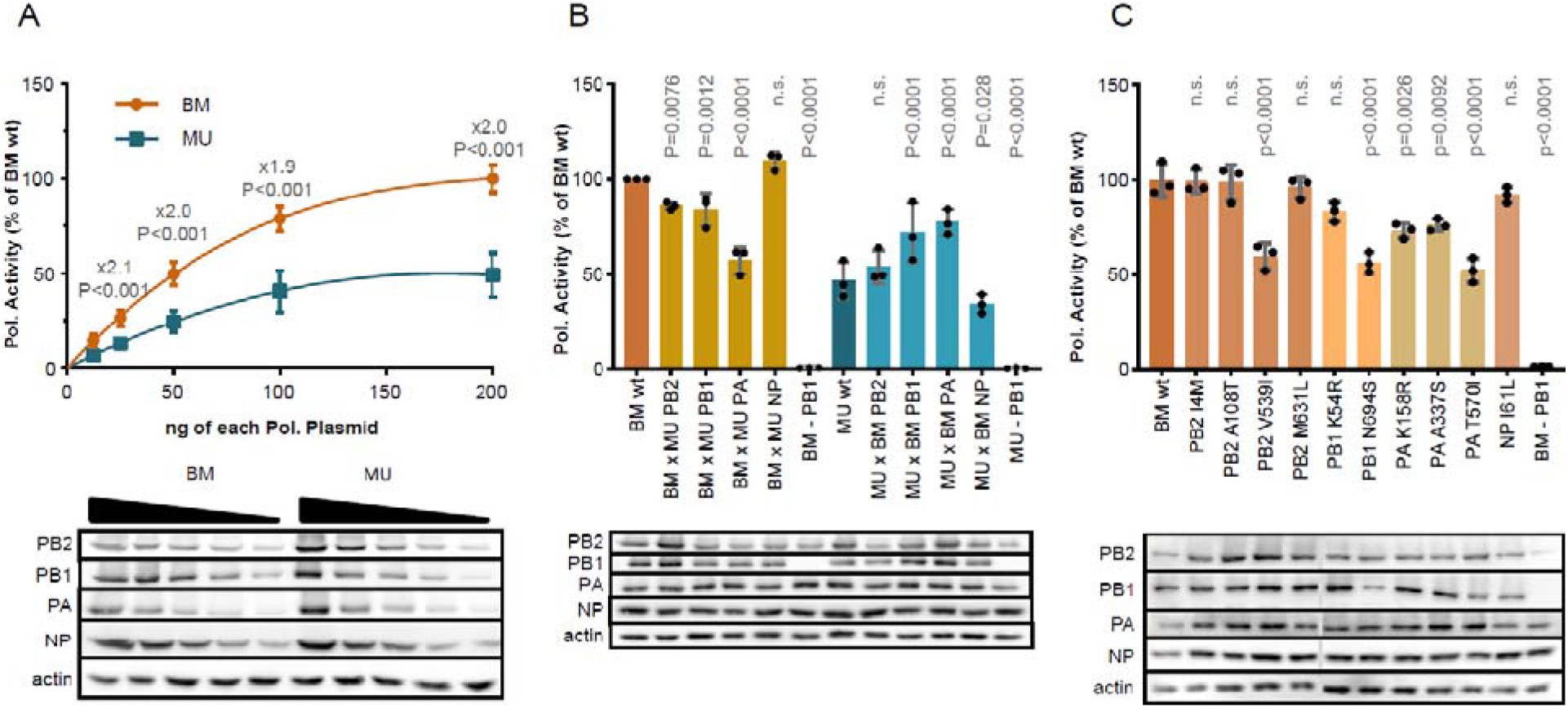
*In vitro* activity of BM and MU-162 polymerases. (A) dose response, (B) effect of whole segment swapping, (C) effect of single BM-to-MU-162 aa changes.

To identify the subunit(s) responsible for this difference, we determined the effect of swapping single polymerase subunits between BM and MU-162 (**Fig. 3B**). The introduction of PA from MU-162 into the BM polymerase complex caused a 1.7-fold reduction of its activity (**Fig. 3B**, P<0.0001). In line with this finding, BM PA increased the activity of the MU-162 polymerase by 1.7-fold (P<0.0001). Exchange of PB1 (1.2-fold reduction for BM x MU-162 PB1, P=0.0012 and 1.5-fold increase for MU-162 x BM PB1, P<0.0001, respectively) and PB2 (1.2-fold decrease for BM x MU-162 PB1, P=0.0076 and no significant change for MU-162 x BM PB1, respectively) subunits caused a similar, but more modest impact on activity. No effect was recorded upon exchange of NP proteins.

To more precisely pinpoint the variations responsible for the observed differences, we generated BM point mutants, each carrying one of the aa changes identified in the MU-162 strain. All three PA mutants significantly reduced the activity of the polymerase compared to BMwt (1.4-fold (P=0.0026) for K158R, 1.3-fold, (P=0.0092) for A337S and 1.9-fold (P<0.0001) for T570I, respectively) (**Fig. 3C**). Of these, K158R was located in the region identified as being of uncertain origin in BM. In PB1, the N694S mutation apparently reduced polymerase activity by 1.8-fold (P<0.0001), which might however be explained by a reduced protein expression level. The V539I aa change in PB2 caused a similar 1.7-fold reduction (P<0.0001) whereas the other three aa changes in this gene did not (including the aforementioned M631L change). These results indicate that, *in vitro*, amino acid polymorphisms encoded in MU-162 affect the activity of the viral RNA polymerase complex. Although this does not necessarily predict functional differences *in vivo*, it is, at this stage, compatible with the notion of phenotypically polymorphic 1918 influenza viruses.

### 1918 influenza viruses and the origin of human seasonal H1N1 influenza viruses

It had long been assumed that seasonal H1N1 IAVs descended (all or in part) from 1918 influenza viruses. The sequencing of the first segments and then genomes of 1918 strains showed their very close phylogenetic relationships with seasonal H1N1 viruses over all segments: non-clock tree reconstructions suggested that the latter directly descended from the former (*17*). However, the development and application of molecular clock models allowing for host-specific rates of nucleotide evolution (thereafter host-specific local clocks; HSLC) supported an alternative scenario, whereby seasonal H1N1 viruses did not evolve directly from the pandemic virus in HA. These models rather suggested that the HA of seasonal IAVs was acquired from a co-circulating homosubtypic H1 IAV through intrasubtype reassortment (*10*). Intriguingly, this scenario is also compatible with other non-genetic information. First, serological studies provided strong evidence that an H3 IAV circulated prior to about 1900 but weak evidence for H3 circulation from ~1900-1918 (*27–29*). Then, the fact that individuals born after 1900 were spared severe outcomes compared to young adults 20-40 years of age was compatible with an initial exposure to H1 viruses better matched to the 1918 pandemic viruses than the putative H3 viruses to which young adults were first exposed to (*10*). We revisited these conflicting hypotheses about the origin of human seasonal influenza using our extended sequence data set and first used the same inference approaches as previously applied. Non-clock maximum likelihood (ML) reconstructions indicated that human seasonal H1N1 and 1918 pandemic viruses cluster together with reasonably high bootstrap support, with the seasonal lineage nested within the 1918 pandemic variants for all segments but PB2 (**Fig. S2**). Time-measured phylogenetic inference using a host-specific local clock (HSLC) model broadly recovers previously reconstructed phylogenetic patterns for all segments but NP. For all segments but HA and NA the human seasonal lineage nested within pandemic flu diversity, while for HA and NA pandemic viruses form a monophyletic cluster with swine influenza viruses, which is positioned as a sister lineage to humans seasonal lineage (*10*) (**Fig. 4a vs b, Fig. S3**); this pattern was previously also observed in NP, but not in our reconstructions.

**Fig. 4:**
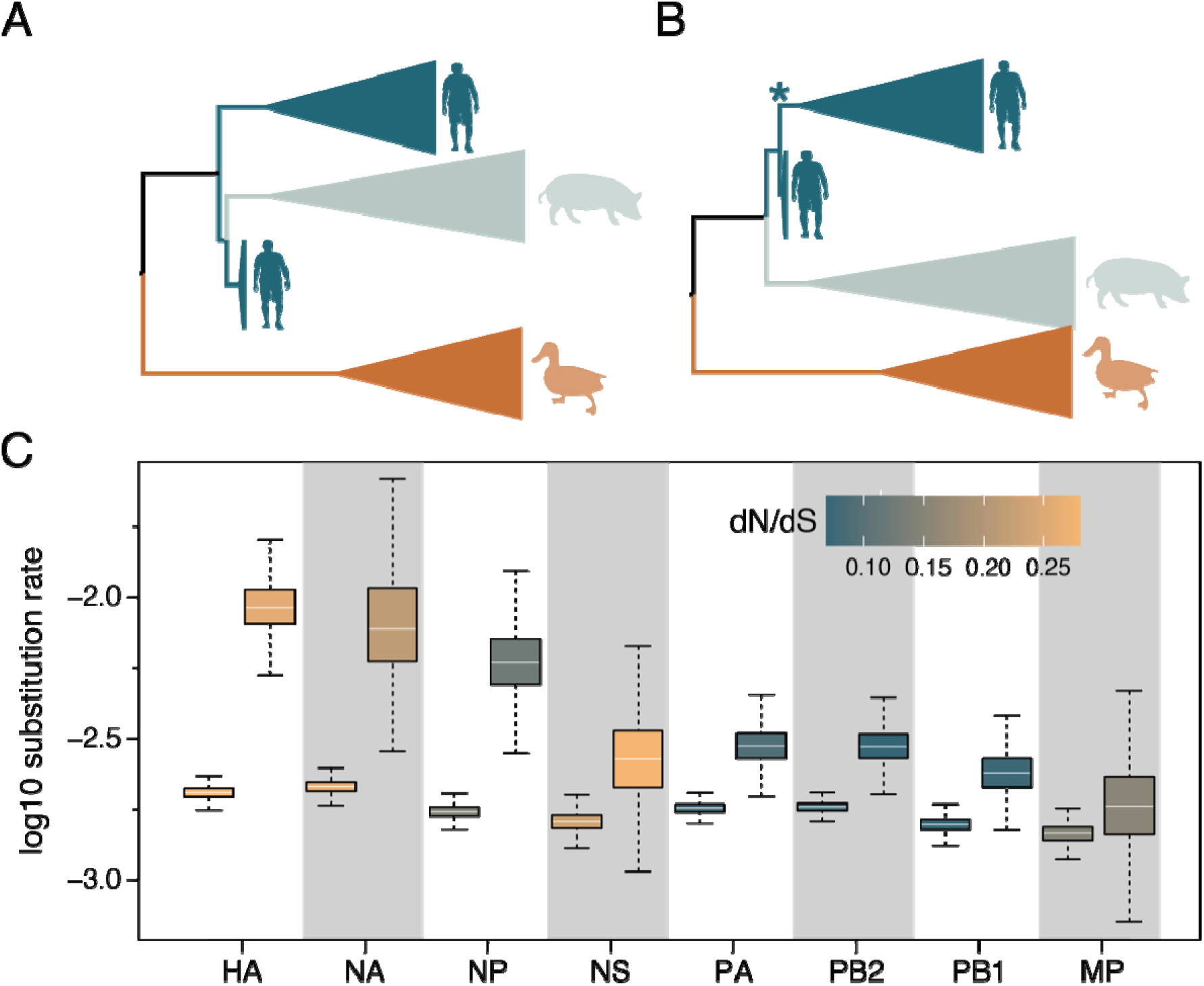
Time-measured phylogenetic patterns and evolutionary rates estimated using Bayesian molecular clock modelling. a) The phylogenetic pattern for HA and NA inferred using a standard host-specific local clock (HSLC) model. b) A monophyletic cluster for human pandemic and seasonal viruses estimated for HA and NA using an extended HSLC model (and using either model for all other segments). The star denotes the branch that is allowed to have a separate evolutionary rate in the HSLCext model. c) Evolutionary rate estimates for the human lineage (first box) and the seasonal ancestral branch (second box) under the extended HSLC model for each segment ordered according to difference in these two rates. The boxes are colored according to the dN/dS estimate for the human lineage and the seasonal ancestral branch in each segment.

Because of the persisting conflict between the difference inference approaches for HA and NA (**Fig. 4a**, **S2 and S3A**), we explored an alternative scenario that allows for a higher evolutionary rate on the branch ancestral to the seasonal lineage. Such a scenario would result in considerably higher divergence between pandemic and seasonal viruses for H1 and N1 than would be expected under a strict clock, and could therefore induce a sister lineage pattern with a relatively deep most recent common ancestor (MRCA) under the HSLC model - which assumes a different but constant rate of evolution in each of the host-specific lineages. We ran simulations that showed that if the rate is elevated on the relevant branch, and this is not accounted for in the clock model, the HSLC inference generally fails to recover the seasonal lineage as a direct descendent of the pandemic virus (**Fig. S5**). Fitting an extended HSLC model that allows for a separate rate on the relevant branch inferred the seasonal lineage as a descendant of the pandemic virus in both HA and NA, consistent with the non-clock trees. Across segments, this model resulted in consistently higher posterior mean rates on the seasonal ancestral branch compared to the seasonal clade rates (**Fig. 4**), but with a variability that largely reflected the variation in selective constraints on the segments. Less stringent purifying selection in HA and NA accompanied by a stronger rate acceleration potentially explains the observed patterns (**Fig. 4c**). To investigate whether variation in selection pressures could explain the acceleration on the branch leading to the seasonal lineage, we performed selection analyses using codon substitution models. These models did not identify diversifying episodic selection or relaxed selective constraints on this branch in any of the segments, implying that only mutation rate/generation time effects may explain the branch-specific elevated rate. Interestingly, a higher mutation rate for the seasonal predecessor (due to a higher replication rate) has indeed been suggested by some experimental evidence comparing the Brevig Mission strain to seasonal strains (*30, 31*).

Altogether, these new analyses revive the scenario of a pure pandemic origin of seasonal H1N1 viruses. However, the essentially phenomenological nature of our modelling approach does not, for now, allow us to definitely favor it over the alternative scenario of a homosubtypic reassortment.

## Conclusion

We derived considerable insight from the addition of only one complete and two partial 1918 flu genomes. Our analyses show significant genomic variation whose spatial distribution suggests frequent long-distance dispersal events, identify potentially adaptive substitutions in NP between first and second wave viruses, hint at viral polymerase phenotypic variation and rekindle the scenario of a pandemic origin for all segments of subsequent seasonal H1N1 influenza viruses. We acknowledge that these findings remain preliminary. Our sample of genomic diversity is extremely small (four good quality genomes) and *in vitro* phenotypic characterization cannot accurately predict *in vivo* phenotypes. Additional genomes from archival samples from various time points starting in 1918, as well as their phenotypic characterization *in vitro* and *in vivo*, will undoubtedly provide the opportunity for more robust tests of our hypotheses. As 3 out of the 4 archival samples from 1918 analyzed yielded good quality viral RNA, influenza genomic research using medical collections can now probably be considered a low risk-high gain enterprise. Therefore, we anticipate that the main obstacle to a better understanding of the evolution of 1918 flu viruses will be the identification of surviving pathological specimens, which highlights the importance of long-neglected museum activities (*32*).

## Author contributions

Conceptualization: LVP, BV, MB, FHL, TS, TW, PL, SCS

Methodology: LVP, BV, MB, MTPG, MAS, MW, TW, PL, SCS

Software: MAS, PL

Validation: LVP, BV, MB, DM, JKT, TS, TW, PL, SCS

Formal analysis: LVP, BV, MB, SL, TS, TW, PL, SCS

Investigation: LVP, BV, MB, AD, JFG, LH, DH, KM, BP, JS, VS, LT, CU, TW, PL, SCS

Resources: FHL, NW, EW, TS, TW, PL, SCS

Data Curation: LVP, BV, MB, AD, JFG, NW, TS, TW, PL, SCS

Writing - Original Draft: LVP, BV, MB, MW, TS, TW, PL, SCS

Writing - Review & Editing: all authors

Visualization: LPV, BV, MB, NW, TW, PL, SCS

Supervision: DH, VS, TS, TW, PL, SCS

Project administration: LVP, SCS

Funding acquisition: FHL, TS, TW, PL, SCS

## Data and materials availability

All raw reads generated for this study have been deposited to the European Nucleotide Archive under project number PRJEB41631 (sample numbers ERS5447401-413). Kraken2 results for all specimens can be visualized through Krona plots at https://zenodo.org/record/4384755. Nucleotide alignments for the 8 viral segments are available at https://zenodo.org/record/4384715.

