## Supplementary Material for "Archival influenza virus genomes from Europe reveal genomic and phenotypic variability during the 1918 pandemic"

**This PDF file includes:**

Materials and Methods

Supplementary Text

Figs. S1 to S12

Tables S1 to S5

#### Materials and Methods

#### Materials and Methods

#### Samples

We obtained 11 lung samples from formalin-fixed specimens identified in the collection of the Berlin Museum of Medical History at the Charité (Berlin, Germany). Sample 447/1913 (sample ID/year of collection) was taken from a 21-year-old (yo) male with a pathological report of fresh confluent pneumonia of the lower lung lobes. Sample 876/1913 was taken from a 45-yo male with a pathological report of calluses on the pleura. Samples 1150/1914 and 928/1915 were taken from a 16-yo and 9½ month-old male, respectively, with a pathological report of tuberculosis. Sample MU-162 was taken from a 17-yo female who died of influenza-related pneumonia in Munich in 1918. The exact collection date was not reported. The pathological finding reported the presence of purulent bronchitis, bronchiolitis and bi-lateral confluent bronchopneumonia. Sample BE-572 was taken from a 18-yo male soldier who died in Berlin on June 27^th^ 1918. The pathological findings were severe purulent pseudomembranous tracheobronchitis and purulent hemorrhagic bronchopneumonia. Sample BE-576 was taken from a 17-yo male soldier who died of influenza in Berlin on June 27^th^ 1918. The pathological findings were fibrinous bronchitis and purulent hemorrhagic bronchopneumonia. Sample 1068/1918 was taken from a 13-yo female. The main pathological finding was fibrinous pseudomembranous bronchitis. Sample 84/1919 was taken from a 55-yo male who died with a caseous pneumonia of the left pulmonary lobe. Sample 247/1919 was taken from a 32-yo woman with tuberculosis who died in Munich. Pathological findings included chronic lung tuberculosis with bronchiectasis, typical lung cavernae and pneumonia. Sample 112/1920 was taken from a 6-month-old individual (gender not reported) who died of influenza. The pathological report included purulent bronchitis and bronchopneumonia, double-sided empyema and collapse of both lungs. Since the exact composition and concentration of the formalin fixative was unknown, we stored the lung samples in PBS to avoid further damage to nucleic acids by adding fresh formalin. Our sample set was further expanded with two formalin-fixed lung specimens preserved within the pathology collection (Narrenturm) of the Natural History Museum in Vienna, Austria, catalogued as influenza-related pneumonia. Sample 18.560/684 originated from the Spital Rudolfstiftung in Vienna and was taken from a child who died on January 6th 1900 with a diagnosis of influenza-related pneumonia. Sample 15.929 originated from the University of Graz and was taken from a 23-yo man who died in 1931. The pathological findings were confluent influenza-related pneumonia with gangrenous cavernae and acute empyema. After sampling, the specimens from the Narrenturm were preserved dry.

Sample preparation

RNA extraction. To maximize chances of viral RNA recovery, we performed 8 separate total nucleic acid extractions from different areas of the lung using the DNeasy® Blood & Tissue Kit (Qiagen) with modifications for formalin-fixed samples, that has been demonstrated to effectively recover both RNA and DNA from such samples (*33*). For each separate extraction, a pea-sized piece (ca. 25mg) of lung was washed in 1 ml PBS to remove residual fixative. The washed tissue was cut into smaller pieces using sterile scissors and added to bead tubes containing tissue lysis buffer (ATL). To reverse formalin-induced crosslinking (*34*), the tissue was heated to 98°C for 15 minutes. To facilitate lysis, the tissue was homogenized by bead beating with a Fast Prep® (MP Biomedicals). We added 20 µl Proteinase K and kept the homogenate at 56°C until the tissue was completely lysed (ca. 1 hour). Subsequent steps were performed according to protocol and nucleic acid was eluted in 35 µl elution buffer (AE).

Library preparation. To maximize viral RNA in the final sequencing libraries, we removed DNA and ribosomal RNA from the nucleic acid extracts before conversion to double-stranded cDNA. For DNase treatment we used the TURBO DNA-free™ Kit (Ambion). To reduce costs for rRNA depletion, 4 DNase-treated extracts were pooled and concentrated using RNA Clean & Concentrator-5 Kit (Zymo Research) and eluted in 13 µl nuclease free water as input for one ribosomal RNA depletion reaction. The other 4 extracts were treated separately. We performed ribosomal RNA depletion and clean-up using the NEBNext® rRNA Depletion Kit (Human/Mouse/Rat) with RNA Sample Purification Beads (New England Biolabs) according to protocol. Following bead clean-up RNA was eluted in 12 µl nuclease free water. We performed cDNA synthesis, using the SuperScript™ IV First-Strand Synthesis System (Invitrogen) and converted cDNA into ds DNA with the NEBNEXT® mRNA Second Strand Synthesis Module (New England Biolabs). Double-stranded DNA was purified using MagSi-NGS^prep^ Plus Beads (Steinbrenner Laborsysteme) and eluted in 50 µl TET. We prepared 5 separate libraries with the NEBNext® Ultra™ II DNA Library Prep Kit for Illumina® (New England Biolabs) without prior fragmentation of double-stranded cDNA and without size-selection upon adapter ligation. All clean-up steps during the library preparation were conducted with MagSi-NGS^prep^ Plus Beads (Steinbrenner Laborsysteme). The libraries were dual indexed with NEBNext® Multiplex Oligos for Illumina® (New England Biolabs), quantified using the KAPA Library Quantification Illumina Universal Kit (Roche), amplified with the KAPA HiFi HotStart ReadyMix (Roche) and Illumina adapter-specific primers, and diluted to a concentration of 4nM for sequencing.

Sequencing. Libraries were sequenced on an Illumina® MiSeq platform using the v3 chemistry (2x300-cycle) and on the Illumina® NextSeq platform using v2 chemistry (2x150-cycle).

#### NGS data analyses

Raw reads were pre-processed using Trimmomatic (*35*), with the following settings: LEADING:30 TRAILING:30 SLIDINGWINDOW:4:40 MINLEN:40. Reads were initially classified using Kraken2 (*36*). Trimmed reads were then mapped to the A/Brevig Mission/1/1918 (BM) reference genome (GenBank accession numbers AY130766, AF333238, AF250356, AF116575, AY744935, DQ208309-11) using BWA-MEM (*37*). Upon detection of influenza reads in sample MU-162, we proceeded with de novo assembly using metaSPAdes (*38*). Scaffolds greater than 750 bp (the smallest BM segment is 830 bp) were blasted against a database of influenza virus sequences and all 8 viral segments were identified. The 8 segments resulting from the de novo assembly were then used as a reference for reference based mapping of reads generated from sample MU-162 and all other samples included in this study. Upon trimming, paired end reads were merged using the Clip&Merge tools from EAGER (*39*). We sorted mapping files and removed duplicates using the SortSam and MarkDuplicates tools from Picard (<http://broadinstitute.github.io/picard>). We assessed RNA damage using mapDamage v2 (*40*). Base calling for MU-162 was set to 20 reads and 95% agreement whereas for BE-572 and BE-576 to 3 reads and 50% agreement. Recombination analyses for the PA segment were run in Rdp4 using the PA dataset assembled by Worobey et al.(*41*) We reduced the dataset to unique sequences using FaBox (*42*) and then chose the 99 most ancient ones to analyse with the MU-162 sequence. We ran Rdp4 (*43*) with all methods (changing the settings of the bootscan approach to a window size of 100). Geneconv, bootscan, lard and 3seq identified Brevig as a recombinant (MU-162 backbone with a ca. 100bp originating in another virus). PhyML (*44*) was finally run on the backbone and recombinant region.

#### Evolutionary analyses and simulations

Datasets. For the HA and NA segments, the newly generated pandemic H1N1 sequence data were complemented with the selection of human, swine and avian H1 and N1 lineages previously used by Worobey et al. (*10*). The A/London/1/1919 isolate (*9*) was also added to the HA data. The newly sequenced samples from Berlin differ from each other in HA at two positions (alignment positions 356 and 1600). BE-576 is polymorphic at these positions with ~40% of the reads exactly matching the BE-572 HA sequence. It can hence not be excluded that BE-576 is identical to BE-572, and that the majority rule consensus base calls at these positions reflect the amplification of potential RNA damages/RT or PCR errors. For this reason, all HA analyses were run in duplicate, once with the majority rule consensus sequence, and once with these positions represented by the appropriate ambiguity code. For the other segments, except for NS, the new 1918 pandemic flu data were complemented with the human H1N1 viruses, the classical swine flu strains and Eastern plus Western avian lineages from the final datasets used by Worobey et al. (*41*) downloaded from Dryad. For NS, all human and swine lineages selected by Worobey et al. (*41*) were used, but only a subset of the avian taxa was taken along to not have to enforce monophyly constraints on the human seasonal and pandemic lineages when accounting for the host-specific evolutionary rates (see below).

Evolutionary analyses. Sequences were aligned with MAFFT v7.313 (*45*) and Aliview v1.19 (*46*) was used for manual refinement of the alignments. Maximum likelihood (ML) trees for all datasets were reconstructed using IQTREE 2.0 (*47*) with a general time-reversible (GTR) nucleotide substitution model (*48*) and a discretized gamma distribution to model among-site rate heterogeneity. Bootstrap support was estimated using the ultra-fast bootstrap procedure with 1000 pseudo-replicates (*49*).

Time-measured phylogenies were estimated using BEAST v1.10 (*16*) with the same substitution model settings as for the ML trees. The skygrid model was used as a flexible demographic tree prior (*50*). To accommodate host-specific rates in the estimation of divergence times, we employed a molecular clock model that allows for different evolutionary rates in lineages of different host species (*51*). The standard host-specific local clock (HSLCstd) model specifies the avian influenza virus rate as the background rate, and assumes a different rate for the human (and swine) lineage, and yet another separate rate for the swine lineage specifically. To keep the model identifiable, we constrain a monophyletic cluster for the human and swine lineage as well as a monophyletic cluster for the swine lineage specifically within this cluster. For this specification it remains unclear which rate to assign to the branch ancestral to the human (and swine) lineage (avian or human) and to the branch ancestral to swine lineage (human or swine). Therefore, we integrate over both alternatives for each branch using a stochastic variable selection procedure (*52*). To reconcile the time-measured phylogenetic reconstructions with the non-clock ML tree reconstructions for the HA and NA segment, we also extend this model to allow for a separate evolutionary rate along the branch ancestral to the human seasonal lineage. We refer to this model as the extended host-specific local clock (HSLCext).We used BEAGLE 3 (*53*) for efficient likelihood computation and simulate the MCMC chains sufficiently long to ensure stationarity and mixing as diagnosed using Tracer v1.7.0 (*54*). We summarized posterior tree distributions in the form of maximum clade credibility (MCC) trees and visualize these trees using FigTree (<http://tree.bio.ed.ac.uk/software/figtree/>). Selection analyses were performed using codon substitution model approaches implemented in HyPhy (*55*). Specifically, we tested for episodic (diversifying) selection and relaxed selection along the branch ancestral to human seasonal viruses in the human H1N1 lineage using aBSREL (*56*) and RELAX (*57*) respectively. For this purpose, we used the human lineage subtrees extracted from the BEAST MCC trees estimated using the HSLCext model.

Simulations. We performed two sets of sequence simulations. First, we simulated sequence data according to the BEAST estimate obtained under the HSLCstd model. We simulated 20 replicate data sets under this scenario (with the same size as the HA data set) and then evaluated the performance of both non-clock maximum likelihood reconstruction and BEAST estimation using the extended HSLC model. Second, we simulated sequence data using the same procedure, but now according to the BEAST estimate obtained under the HSLCext model and evaluated the performance of both non-clock maximum likelihood reconstruction and BEAST estimation using the standard HSLC model. Simulations were performed using piBUSS (*58*) .

#### Functional analyses

Plasmids. Complete cDNAs of influenza virus genomic segments derived from sample MU-162 or of strain A/Brevig Mission/1/1918 (H1N1, BM) were commercially synthesised (Biocat, Heidelberg, Germany for MU-162 and Geneart, Regensburg, Germany for BM, respectively). Point mutations were introduced into BM segments with the Quikchange II Site-directed mutagenesis kit (Agilent, Santa Clara, CA, USA) according to manufacturer’s instructions. Coding sequences of polymerase and NP proteins were amplified by PCR with the Phusion Green Hot Start II High Fidelity DNA polymerase (ThermoFisher). PCR products for PB2, PA and NP were cloned into the BsaI sites of vector pCAGGS∆Bsa-Blue (*59*). PB1, which contains internal BsaI sites, was cloned into the BsmBI sites of a modified version of the pCAGGS plasmid. All constructs were confirmed by Sanger sequencing. Primers used for cloning and mutagenesis are listed in table S1.

Polymerase activity assay. Trypsinized 293T cell cultures were transiently transfected in suspension in 12 wells using lipofectamine 2000 (Invitrogen) with 50 ng of each pCAGGS-pol plasmid, 100 ng pCAGGS-NP plasmid, and 125 ng of pPolI-NS-Luc, expressing an influenza virus-like RNA encoding a firefly luciferase (*60*). Transfection was normalized by the constitutively expressed Renilla luciferase encoded on plasmid pTK-RL (10 ng; Promega, Madison, WI, USA). For titration experiments, a range from 12.5 to 200 ng of each pCAGGS-pol construct was used, with respective double amounts of corresponding NP plasmid, while luciferase plasmids were kept constant. Total DNA amount was equalized in every sample with pCAGGS. At 24 h post transfection, luciferase activity was measured with the Dual-Luciferase® Reporter Assays System (Promega, Madison, WI, USA) on a Tristar LB 941 luminometer (Berthold Technologies, Bad Wildbad, Germany) according to manufacturers’ instructions.

Immunoblotting. Equal amounts of cell lysates from luciferase assay replicates were pooled, denatured in reducing SDS-PAGE sample buffer for 5 min at 95°C, separated on 8% SDS gels, subjected to immunoblotting with antibodies against PB2 (rabbit, ThermoFisher/Pierce, #PAS-32221), PB1 (rabbit, ThermoFisher/Pierce, #PAS34914), PA (rabbit, GeneTex, #125932), NP (mouse, Acris, #AM00929PU-N) and actin (mouse, Sigma, #A2228), respectively and detected with HRP-labelled anti-mouse (DAKO, #P0260) or anti-rabbit antibodies (DAKO, #P0217), respectively.

Statistical analysis. All values are expressed as mean, error bars indicate standard deviations. Sample sizes, that is, number of biologically independent experiments, are indicated in figure legends. Each biological experiment contained three technical replicate values. For multiple comparisons of several groups of a single variable (polymerase subunit exchange, mutant analysis), nested one-way ANOVA was used and P-values were corrected for multiple testing with Sidak’s method. For multiple comparisons of three or more groups with two independent variables (titration of polymerase activity as a function of plasmid concentration), nested two-way ANOVA was used and P-values were corrected for multiple testing with Sidak’s method. Curve fitting was done with regression analysis using 3^rd^ order polynominal equations. Calculations were performed with GraphPad Prism, version 8.1.2.

#### Human RNA preservation screening

The influenza positive samples were analysed for human RNA preservation. The sequencing data was processed using EAGER (*39*). Sequencing reads were inspected with FastQC before merging and adapter trimming with AdapterRemoval V2.2.1a. Pre-processed reads were then aligned to the human transcriptome reference (Gencode, Release 34), using BWA with a minimum quality score of 0 and a maximum edit distance of 0.01. The MarkDuplicates method was used for duplicate processing (Broad Institute) as well as DamageProfiler v0.3.8 and Qualimap (*61*) to investigate the read length distribution and the damage patterns (Fig. S9-11).

### Supplementary text

#### S1. Histopathology

We performed histopathological analyses of the MU-162, BE-572 and BE-576 lung specimens. For conventional histopathology, slides of lung tissue were stained with hematoxylin and eosin. A gram staining was also performed to identify gram-positive bacteria. Sample MU-162 was characterized by severe purulent, partially confluent bronchopneumonia with small hemorrhages and edema without clear histomorphological evidence of bacterial colonization (fig. S1A and S1B). Sample BE-572 was characterized by severe purulent, partially hemorrhagic bronchopneumonia with alveolar edema and evidence of bacterial colonization by gram-positive cocci (fig. S1C and S1D). Sample BE-576 was characterized by severe pulmonary alveolar edema with pronounced hemorrhages, emphysema and first stages of acute bronchopneumonia with evidence of bacterial colonization by gram-positive cocci (fig. S1E and S1F).

#### S2. Evaluation of human RNA preservation

Fully formed RNA virus particles are built to protect the integrity of the viral genome under various conditions for days or even months (*62*). This might also affect RNA recovery from formalin-fixed tissue, due to a better RNA protection prior to fixation in comparison to the endogenous RNA. Hence, we assessed the endogenous human RNA preservation by mapping all reads to a full human transcriptome reference (table S2). Strikingly, most of these fragments show a significantly lower average length than the fragments mapping to the IAV reference (table S5). The smaller fragments of human endogenous RNA compared to the viral RNA are a first indication of the protective function of the viral capsid in RNA recovery from formalin-fixed wet specimens. The length of fragments mapping only to the transcriptome varied greatly between the samples and within independent extractions from the same sample. This might be due to varying time between patient death and formalin fixation, the exact formalin composition (e.g. buffered or unbuffered formalin and formalin concentration), different penetration of the fixative in different parts of the tissue, or remaining gDNA fragments which leads to RNA fragmentation during the RNase H treatment of the rRNA depletion step.

S3. Reconciling clock and non-clock topologies

Non-clock maximum likelihood phylogenetic reconstruction indicates that human seasonal H1N1 and 1918 pandemic viruses cluster together with reasonably high bootstrap support, with the seasonal lineage nested within the 1918 pandemic variants, for both the HA and NA segments (fig. S2), while inference under the standard HSLC model places the human seasonal lineage as a sister clade of the classical swine flu and 1918 pandemic lineages (fig. S3).

If the human seasonal and pandemic lineages are indeed monophyletic, the incompatible pattern under the standard HSLC model - which assumes a constant rate of evolution in each of the host-specific lineages - could be induced by a considerably higher rate of evolution in the years following the pandemic. This would result in a considerably higher divergence between pandemic and seasonal viruses for H1 and N1 than expected under a strict clock, and could therefore induce a sister lineage pattern with a relatively deep MRCA in the time-measured reconstructions. To explore this hypothesis, phylogenies including only the human H1 and N1 taxa were estimated using the same substitution and demographic models as under the standard HSLC but specifying a relaxed clock model (*63*).

For HA, this indicates an elevated evolutionary rate on the branch ancestral to the human seasonal MRCA, and results in a topology that is compatible with the non-clock ML phylogeny (fig. S4). For NA, a pattern similar to that of HA emerges (fig. S4). These results are in line with the idea that significant divergence that accumulated after 1918-1919 could indeed induce the sister pattern under a constant rate assumption over the entire lineage.

S4. Simulation-based assessment of standard and extended host-specific local clock model inference and non-clock phylogenetic inference.

The simulations under the extended host-specific local clock (HSLCext) pattern (fig. S5), as estimated for the HA segment, indicate that the non-clock ML tree reconstructions correctly infer a monophyletic clade for the 1918 pandemic and human seasonal virus in 17 out 20 cases, and incorrectly infer a deep TMRCA in 3 out of 20 replicates (bottom two panels). Not accommodating the higher rate on the branch ancestral to human seasonal results in a biased inference of a deep TMRCA in 17 out of 20 replicates for the standard HSLC (HSLCstd) model.

The simulations under the HSLCstd pattern (fig. S6), as estimated for the HA segment, indicate that both the non-clock ML tree reconstruction as well as the HSLCext model, which allows for a different rate on the branch ancestral to human seasonal, are able to recover the right topology with a 0.95 probability. In line with this, the extended HSLC model does not infer a significantly positive effect on the branch ancestral to human seasonal (not shown). This indicates that the extended HSLC model and the non-clock ML inference are unlikely to produce biased results.

S5. HSLC standard versus HSLC extended: impact on clustering of human seasonal H1N1

For each segment, the plausibility that the seasonal lineage directly emerged from the pandemic diversity increases under the extended HSLC model as compared to under the standard HSLC model (fig. S3 and Table S3). This is particularly outspoken for the HA segment: where the results under the standard HSLC model are in line with reassortment between the human pandemic lineage and an antigenically distinct co-circulating H1 virus (*10*), the clustering patterns under the extended HSCL model suggest otherwise. Support for different evolutionary trajectories of the human seasonal lineage under the HSLCext model varies depending on the inclusion of short HA sequence fragments of pandemic isolates, pointing to a need for more (near) full length HA sequences from around the time of the pandemic. For NP and PA, we find perfect support for nested clustering of the seasonal taxa within the diversity of their pandemic counterparts under both clock models. Support for this is high to perfect for MP, NS and PB2, and is low to moderate for NA and PB1.

S6. Evidence for multiple transatlantic migrations

Analyses of the HA with and without ambiguity at alignment positions 356 and 1600 show that the A/London/1/1918 strain, sampled on 1918-11-13 and representing the first wave of the epidemic in the 1918-1919 winter in London, is most likely not from the same pandemic lineage that caused the second wave of the epidemic that winter (represented by A/London/1/1919, sampling date: 1919-02-15) (.75 and .73 PP, Fig. 2 and fig. S7). This view is reinforced by a separate analysis of the 1918 HA clade (for which we assumed an exponential growth model and a strict clock): posterior support for different epidemic lineages in the first and second wave in London in the 1918-1919 winter increases to .89 and .87. Like the London samples, the isolates from Germany cluster in accessbetween pandemic variants sampled in North America (fig. S7 and S8). This is in line with extensive transatlantic mixing of pandemic influenza. To more explicitly assess support for a migration scenario, the relative fit of models in which the clustering of the European isolates was not constrained versus constrained to be monophyletic was determined using the Generalised Stepping Stone sampling. For this, the HA 1918 clade was analysed separately as before. In line with the clustering patterns (Fig. S8), the unconstrained model clearly better fits the pandemic clade, irrespective of the use of ambiguity characters in the BE-576 sequence (ln BF 8.7 and 9.4).

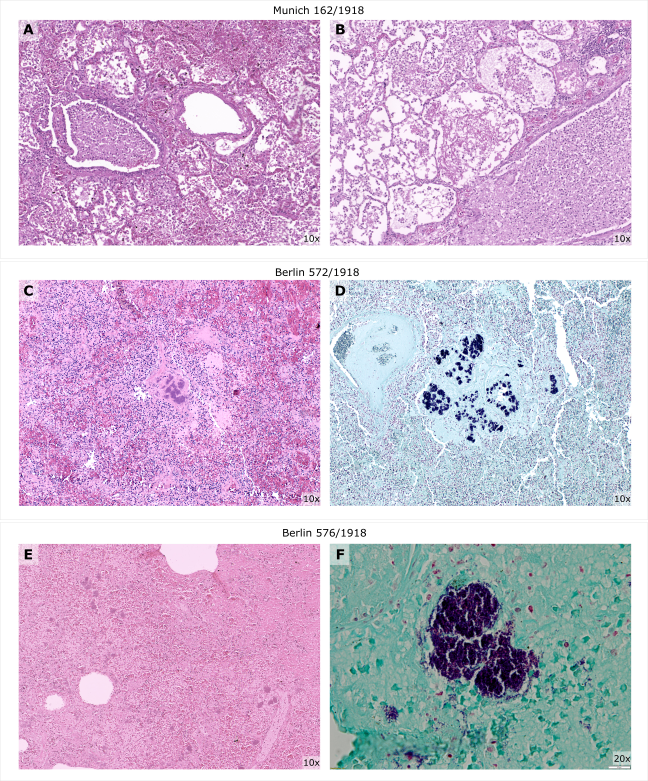

Fig. S1. Histopathological findings of influenza-positive lung specimens. A and B display H&E staining of sample MU-162, C and D show H&E and gram staining for sample BE-572, respectively, E and F show H&E and gram staining for sample BE-576, respectively. Images were taken using either a 10X or 20X objective, which corresponds to a 100X or 200X magnification, respectively.

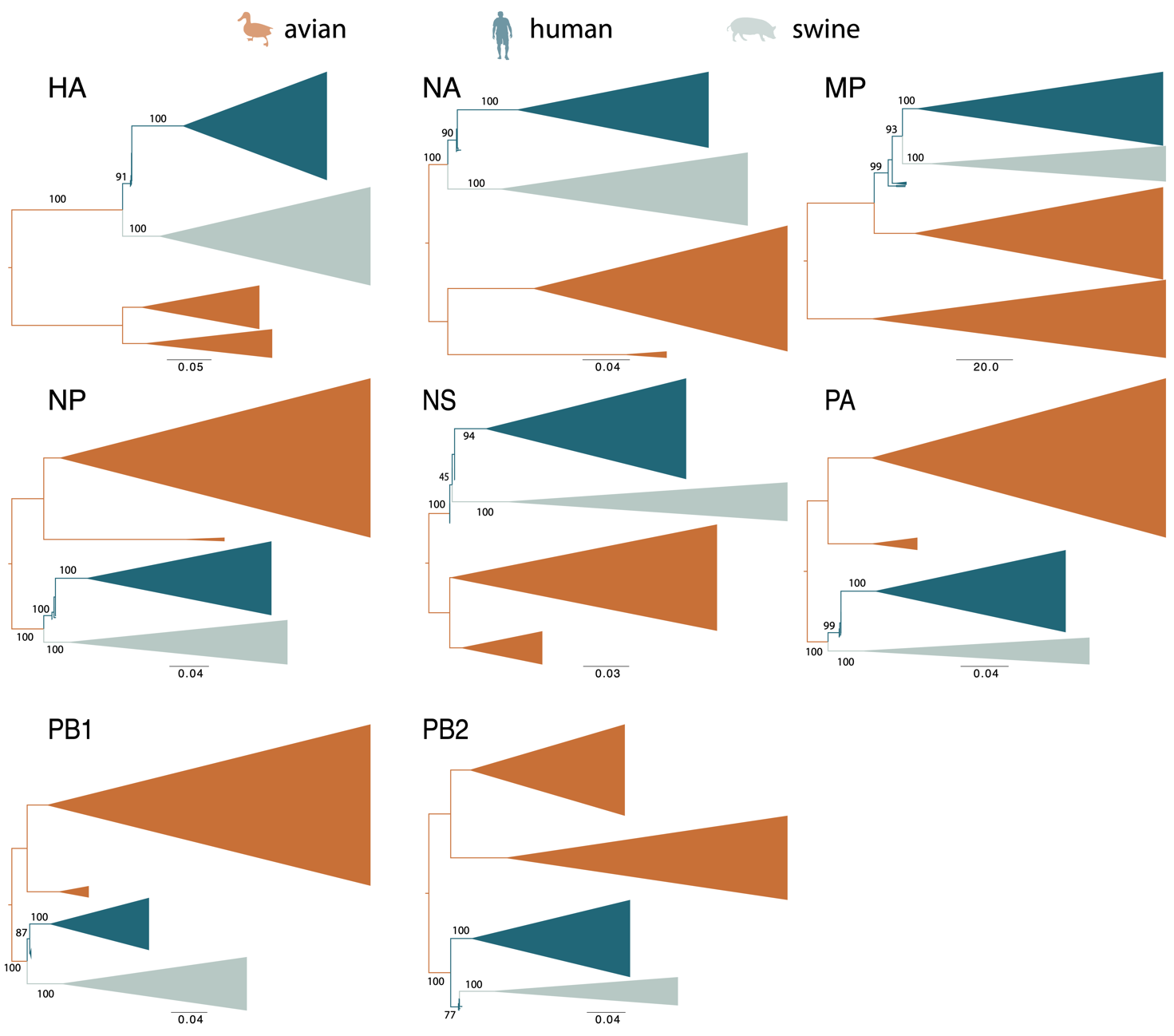
**Fig. S2. Maximum likelihood phylogenetic reconstructions for all segments.** Lineages are colored according to host and numbers at the major nodes of interest represent the percentage bootstrap support.

**A.**
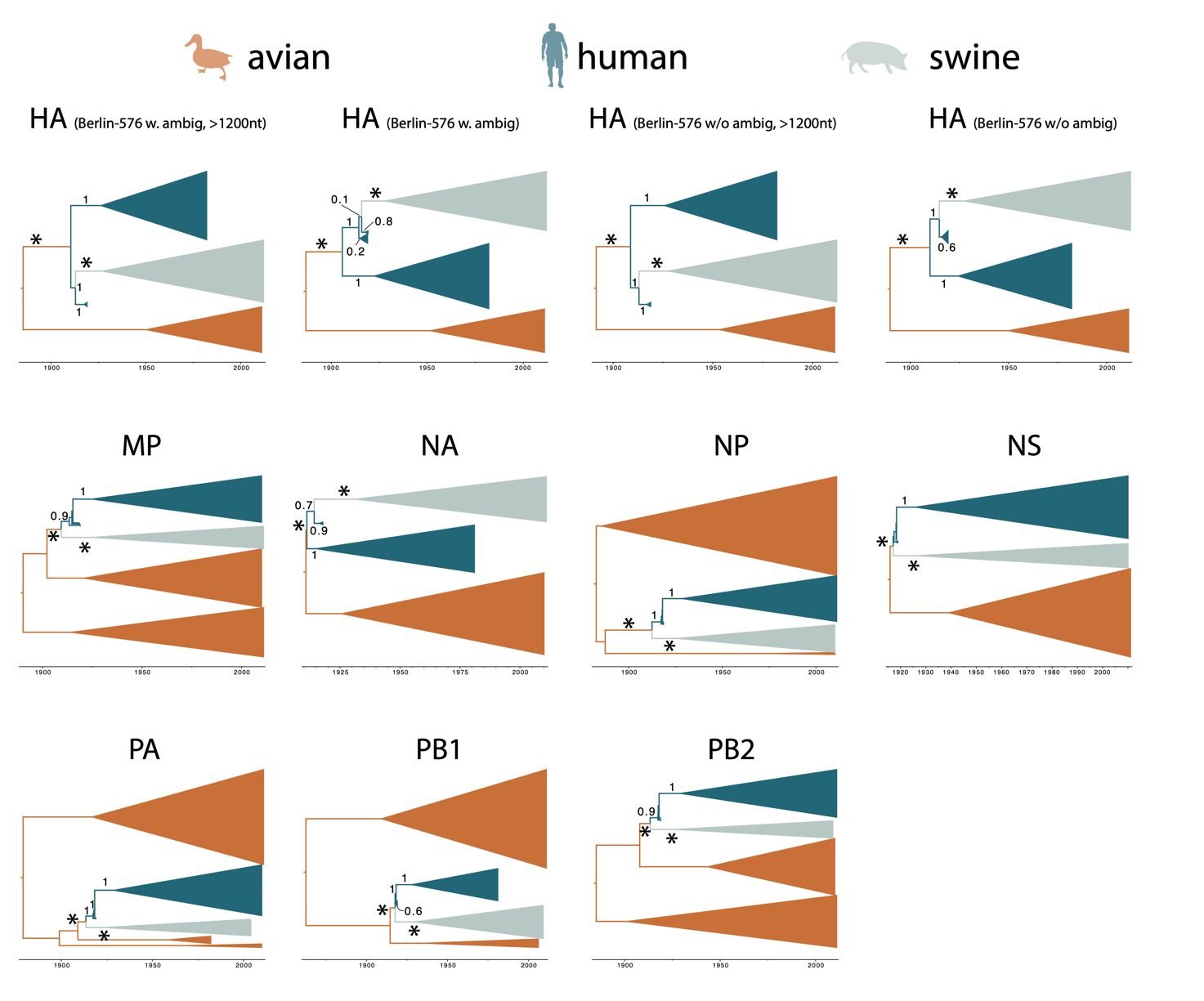

**B.**

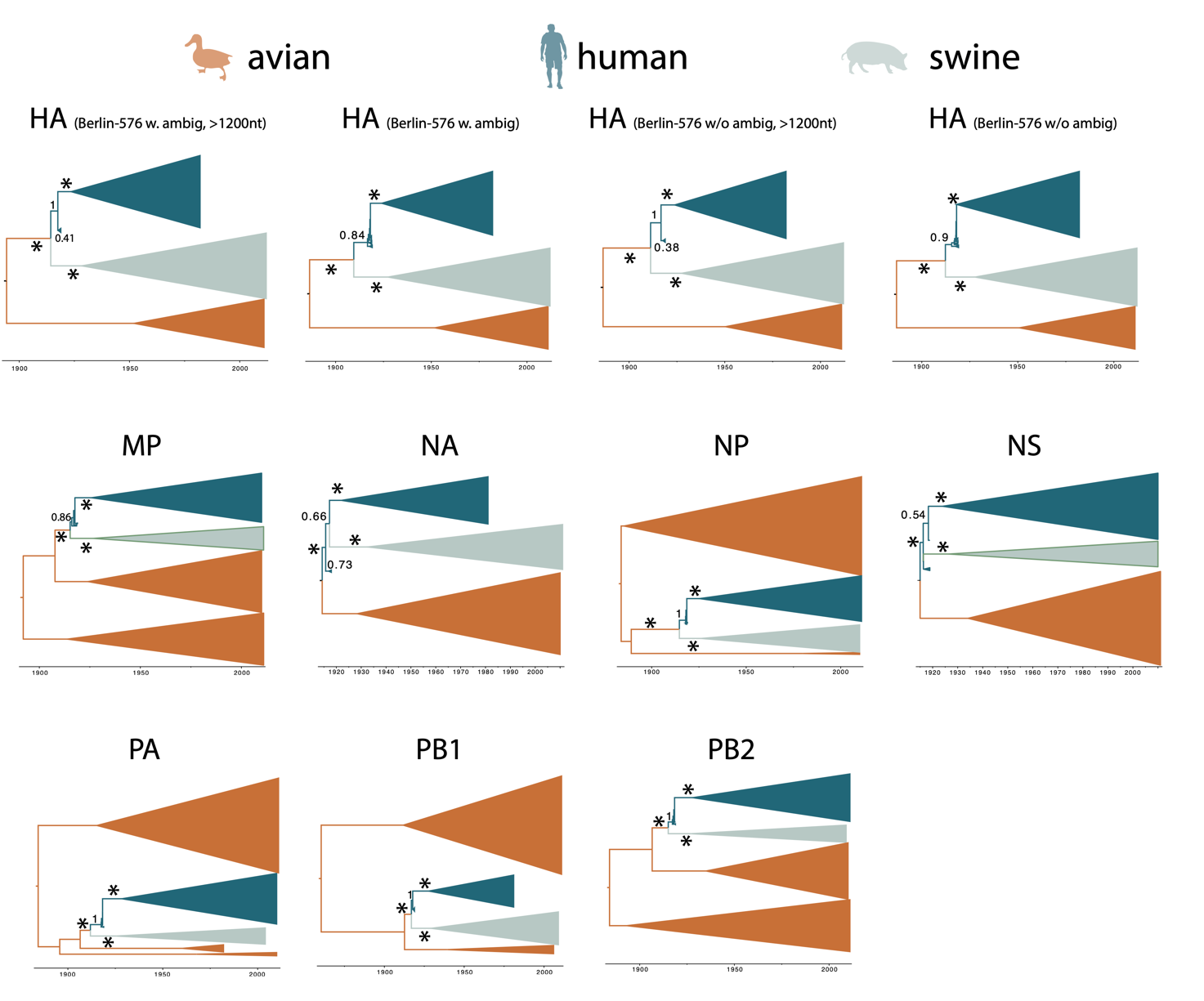

**Fig. S3.** **Illustration of the rate category specification. The trees represent maximum clade credibility (MCC) summary trees for each segment.** The correspondence between branch colors and host category is as in the legend. Asterisks indicate nodes for which a monophyly constraint was specified in order to maintain identifiability of the model. Numbers next to branches refer to their posterior support. Panel A shows the MCC trees inferred under the standard HSLC model, and panel B shows the MCC trees inferred under the extended HSLC model. For HA, '>1200nt' indicates that only human pandemic taxa of length > 1200 nt were considered.

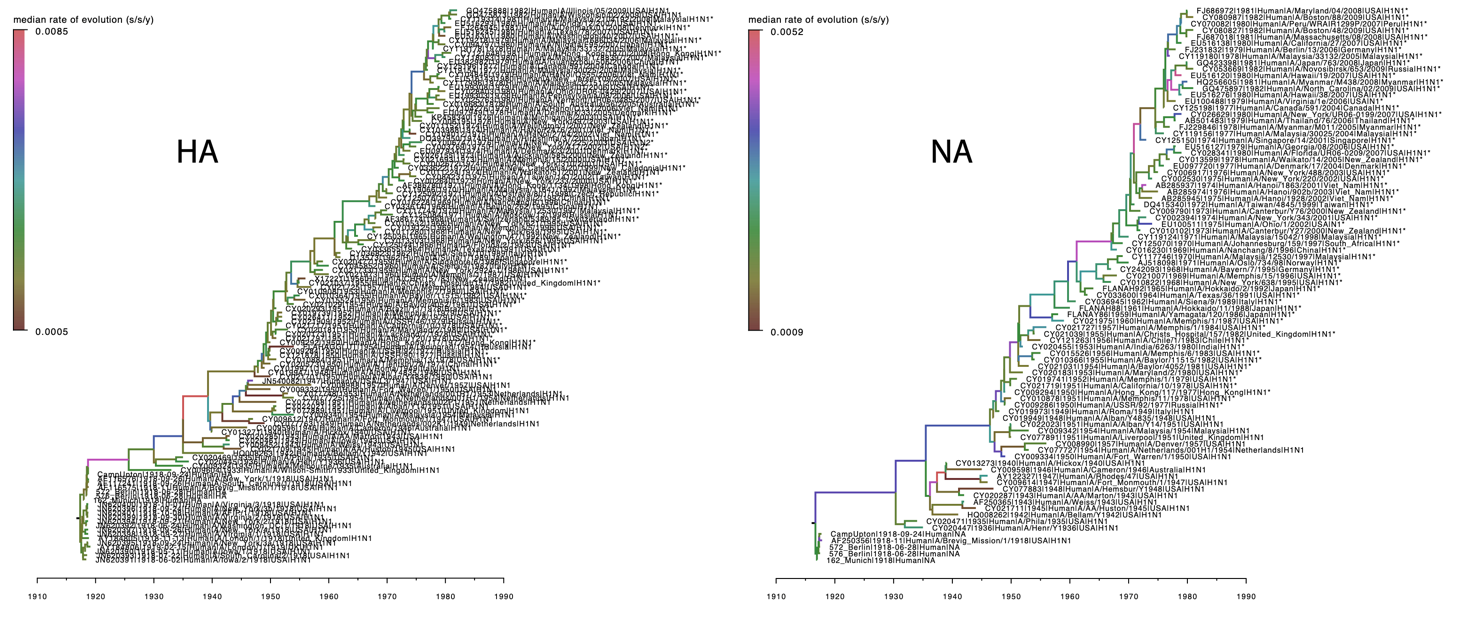
**Fig. S4**. MCC summary trees of human H1 and N1 inferred under a relaxed clock model. Branches are colored according to the median of the rate of evolution estimated over that branch. Correspondence between branch colors and the rate of evolution is as in the legend. For HA, the BE-576 genome with ambiguities was used.

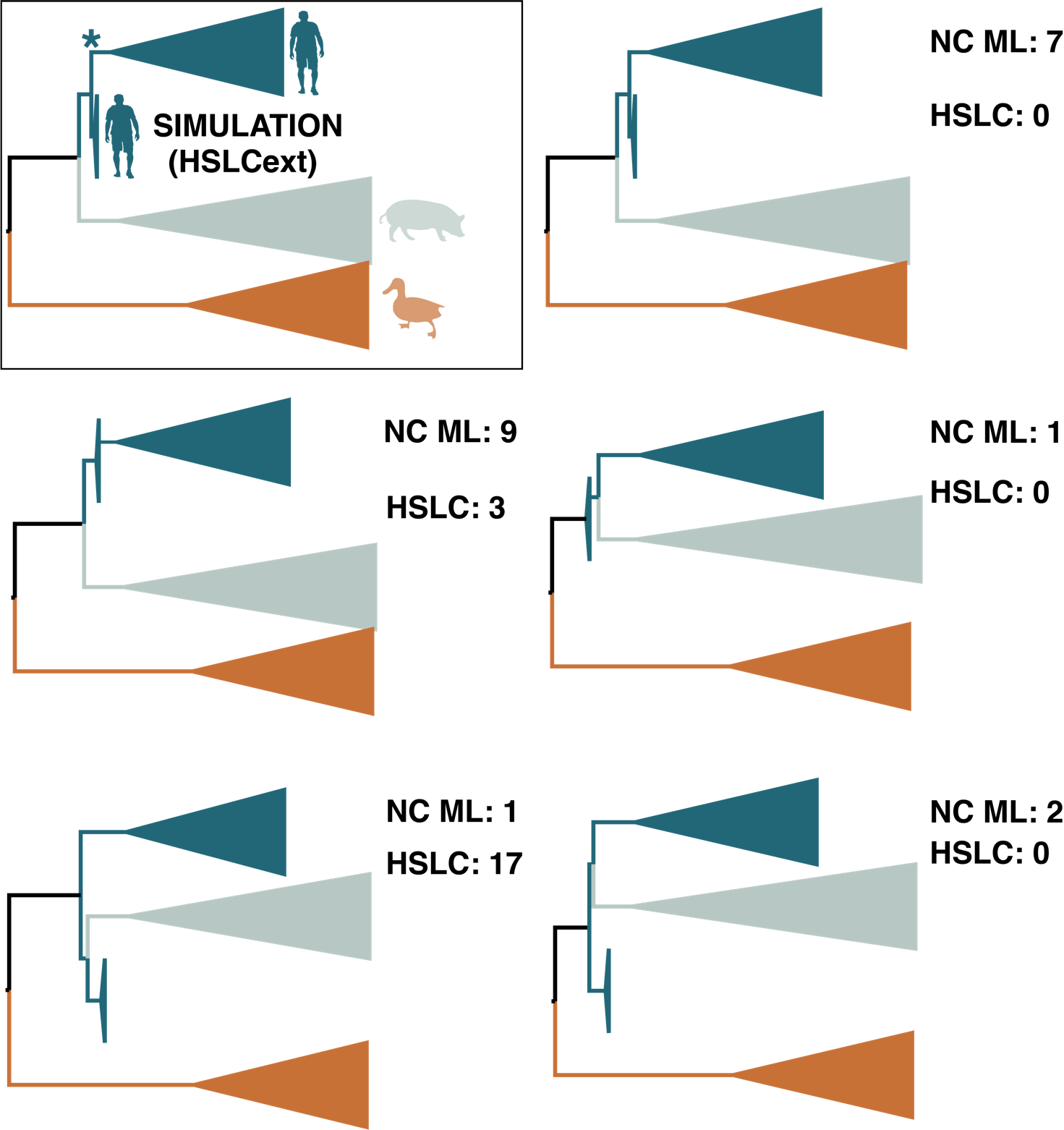

**Fig. S5. Simulations under the extended host-specific local clock (HSLCext) model estimate for the HA segment.** The top left time-measured phylogenetic tree with a grey background represents the estimate under the HSLCext model for HA. A star denotes the branch that is allowed to have a separate rate relative to the standard HSLC model (HSLCstd). 20 data sets were simulated under this scenario and both non-clock maximum likelihood (NC ML) phylogenetic estimation and BEAST estimation using the HSLCstd model were performed on the replicate data. Five topologies are used to summarize the estimates for both methods.

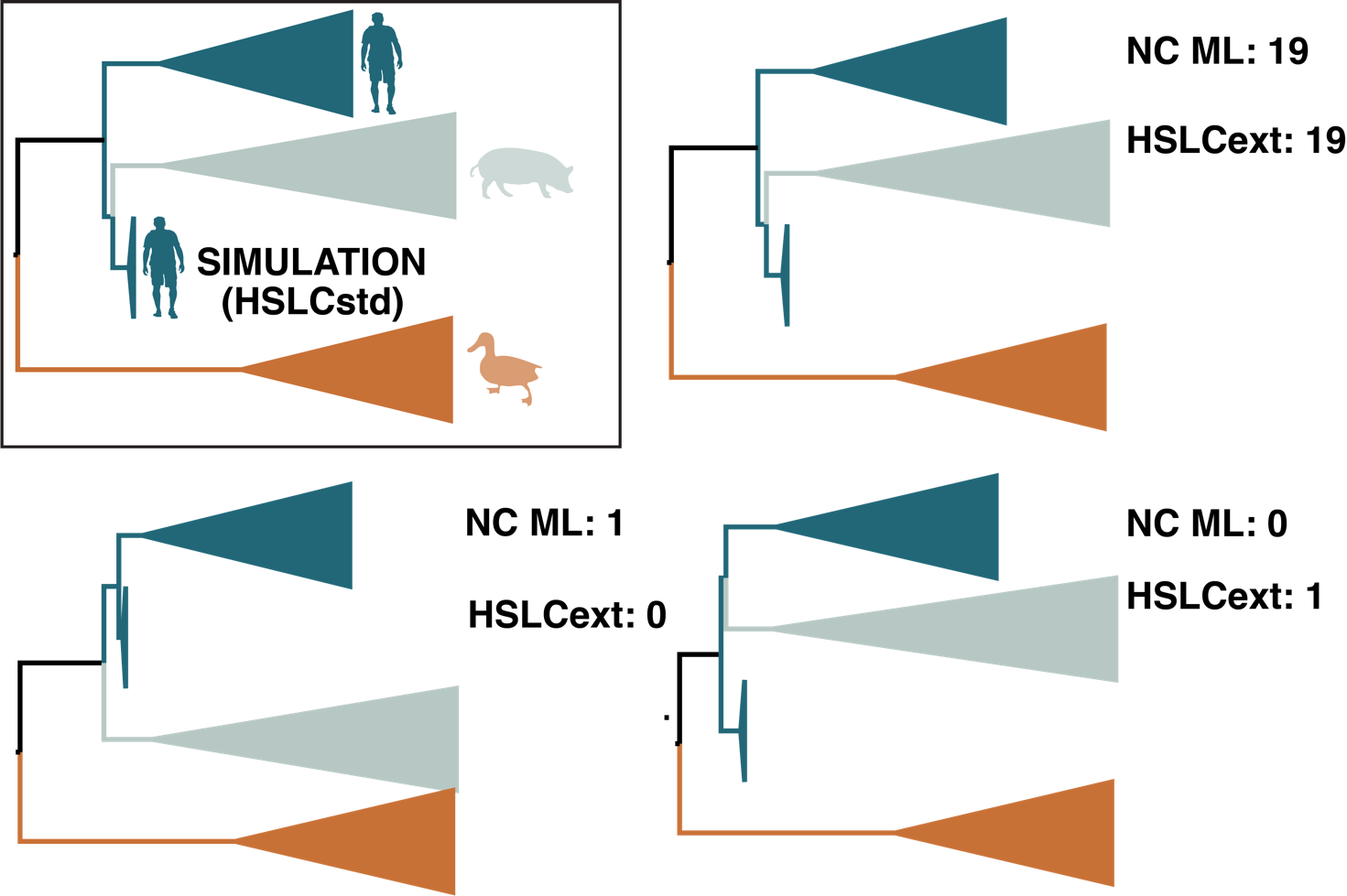

#### Fig. S6. Simulations under the standard host-specific local clock (HSLCstd) model estimate for the HA segment. The top left time-measured phylogenetic tree in a black box represents the estimate under the standard HSLC model for HA. 20 data sets were simulated under this scenario and both non-clock maximum likelihood (NC ML) phylogenetic estimation and BEAST estimation using the extended HSLC (HSLCext) model were performed on the replicate data. Three topologies are used to summarize the estimates for both methods.

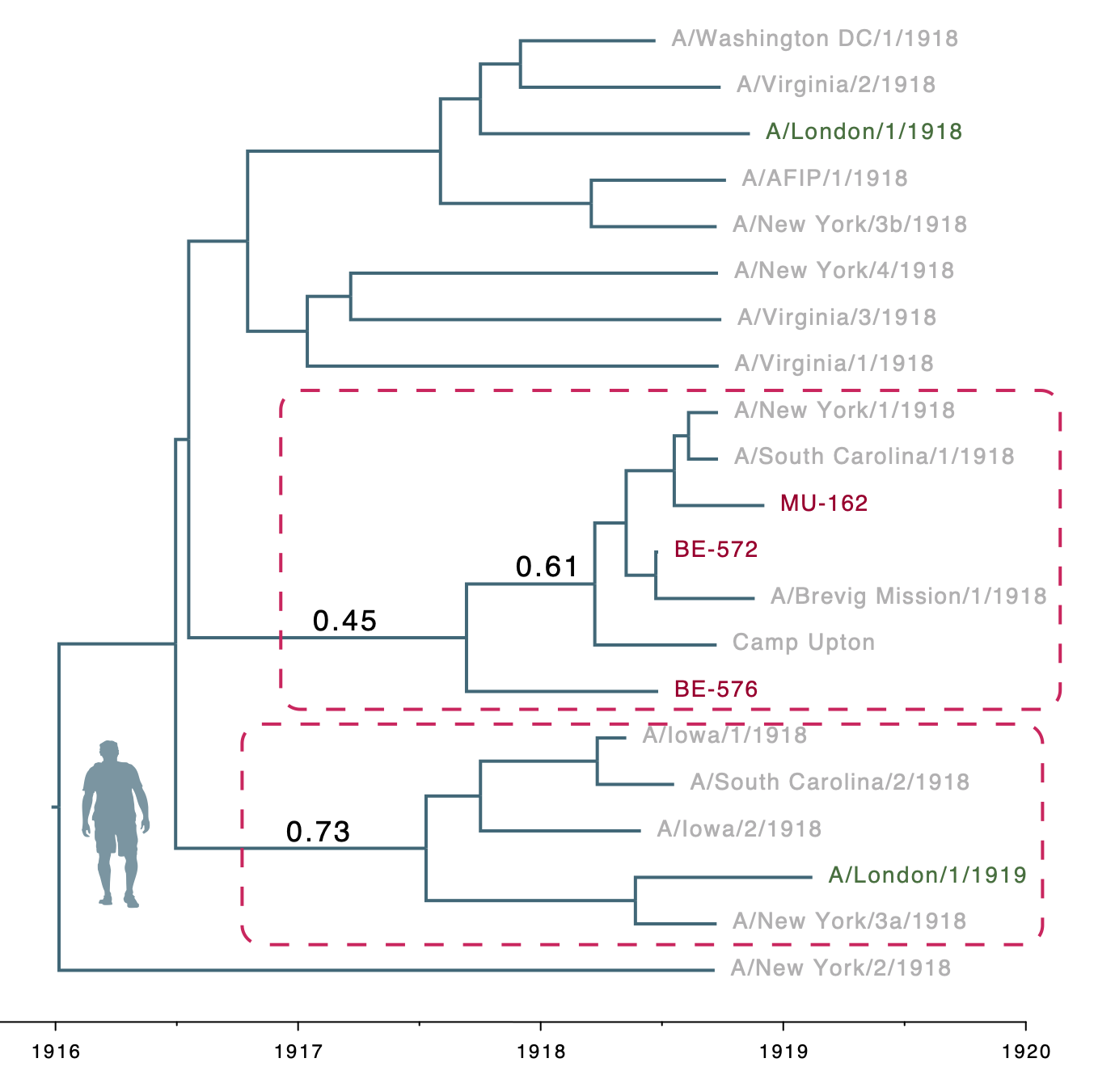
**Fig. S7. Focus on the H1 human pandemic clade of HA using the majority rule consensus for BE-576.** To facilitate identifying the intermixing of the London, German and Northern American lineages, the tip names are colored dark green, dark red and light grey, respectively. Numbers next to branches refer to their posterior probability, which can be interpreted as the probability of the clade being true given the data, the model, and the parameter priors. Red dashed rectangles highlight the clades of interest.

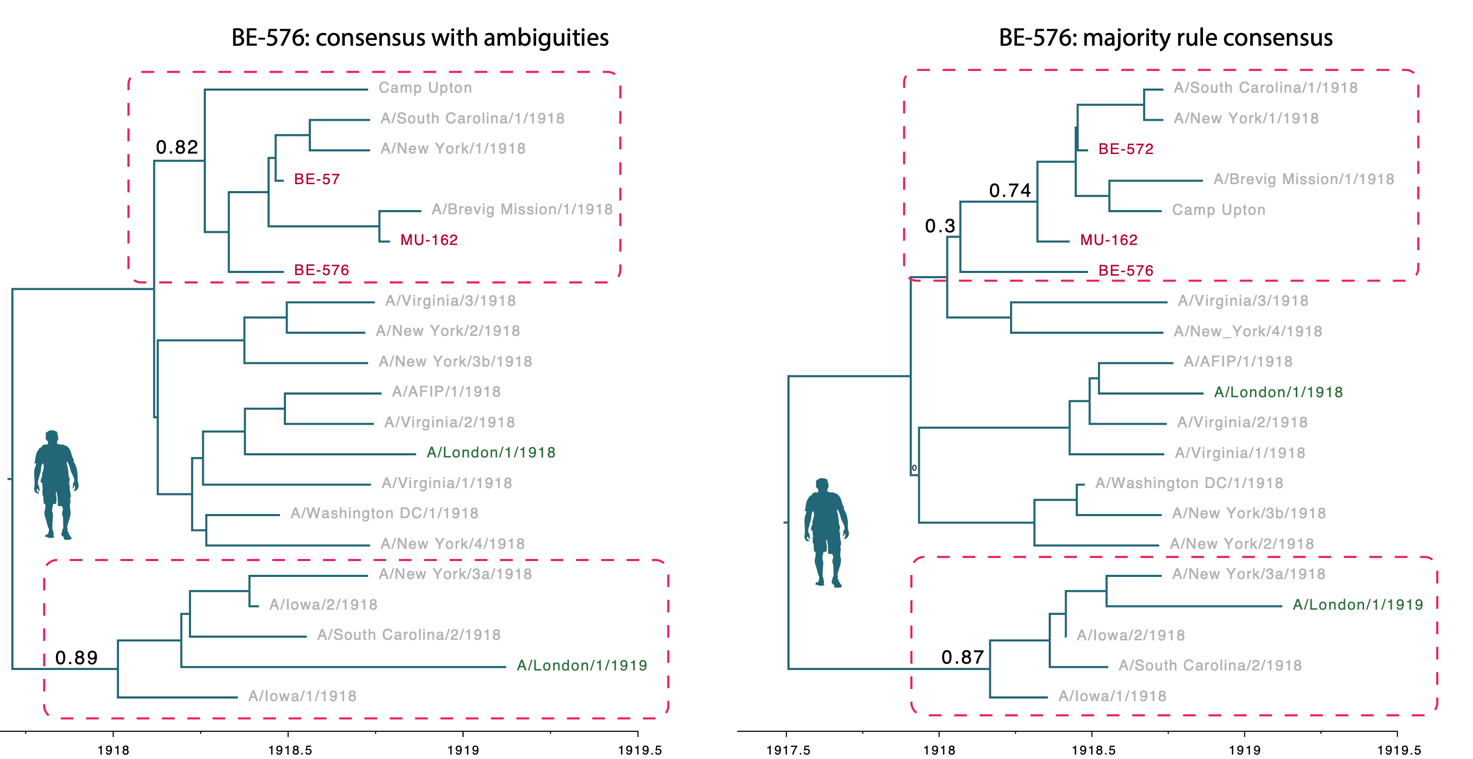
**Fig. S8. The H1 human pandemic clade estimated only from the pandemic HA sequences.** To facilitate identifying the intermixing of the London, German and Northern American lineages, the tip names are colored dark green, dark red and light grey, respectively. Numbers next to branches refer to their posterior probability, which can be interpreted as the probability of the clade being true given the data, the model, and the parameter priors. Red dashed rectangles highlight the clades of interest.

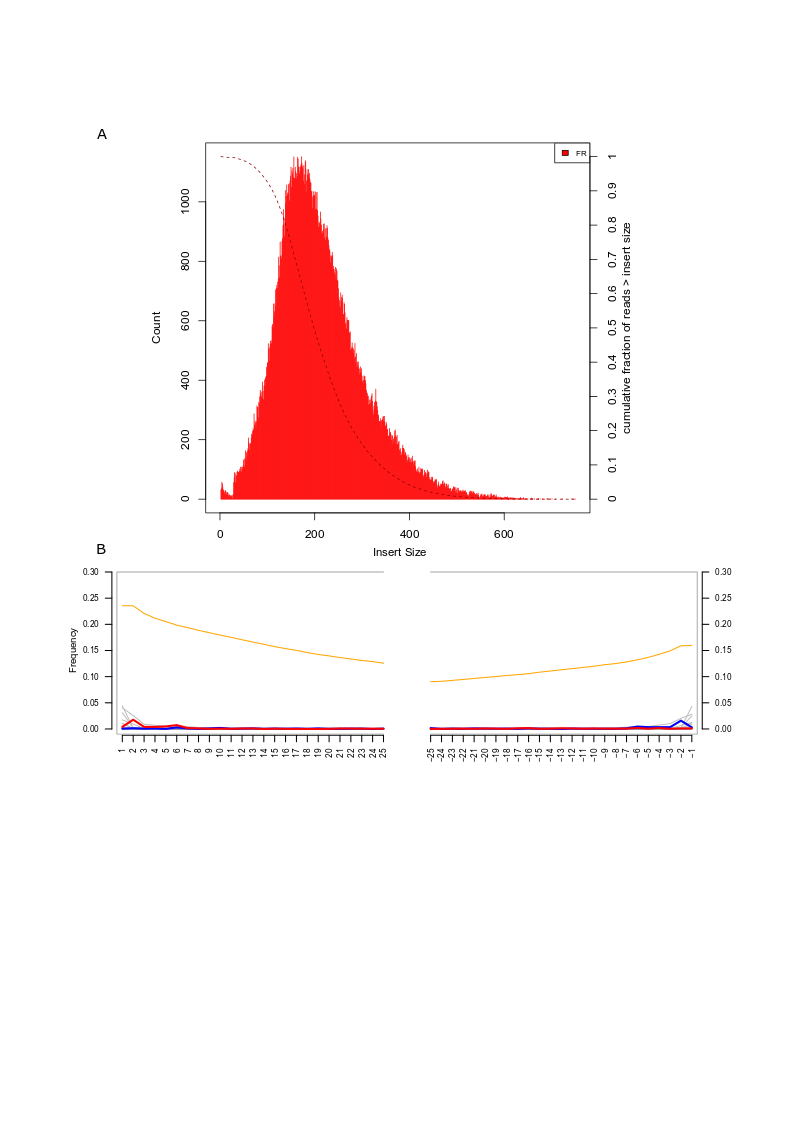

#### Fig. S9. Insert size distribution (A) and mapDamage profile (B) of reads in library 162_9 (MU-162).

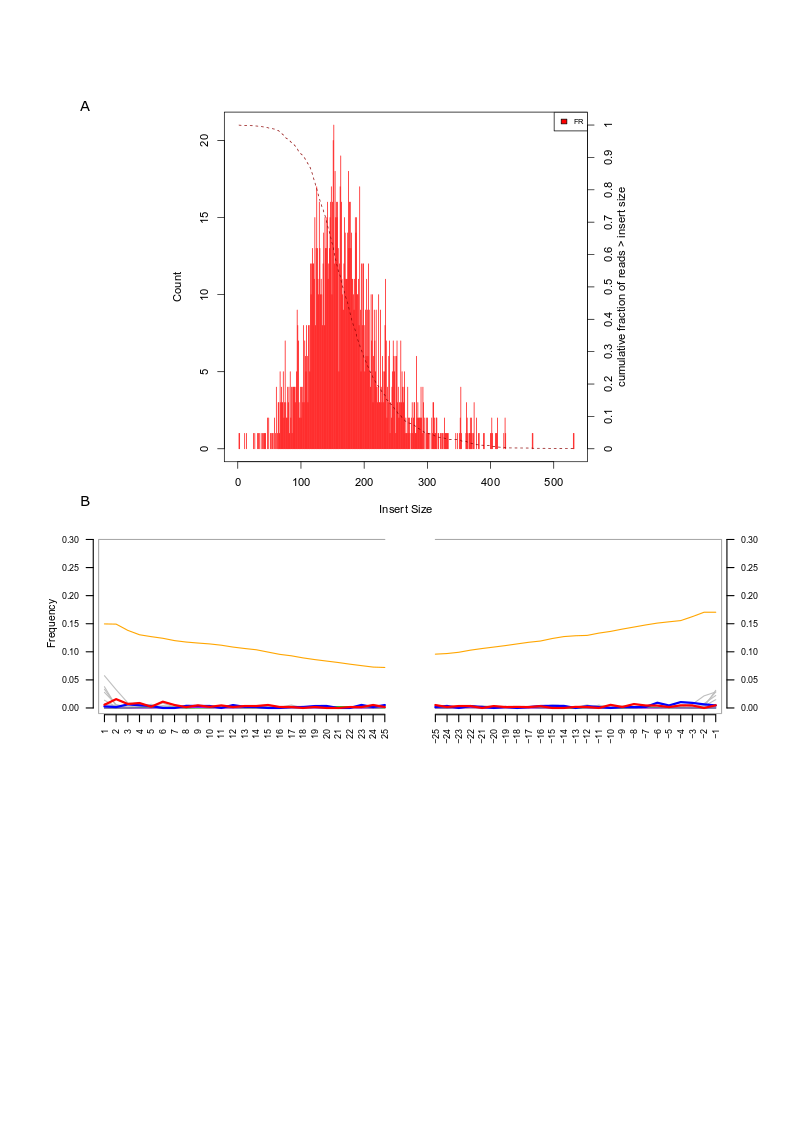

**Fig. S10. Insert size distribution (A) and mapDamage profile (B) of reads in library 572_1 (BE-572).**

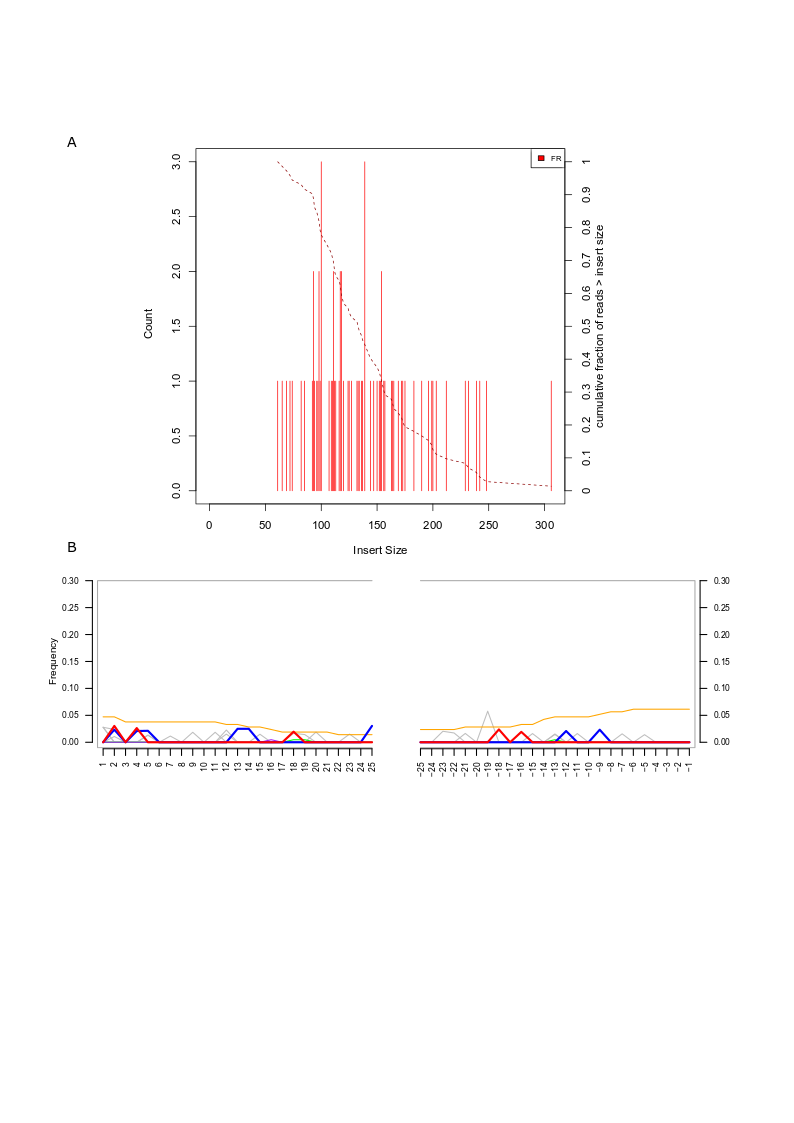

**Fig. S11. Insert size distribution (A) and mapDamage profile (B) of reads in library 576_1 (BE-576).**

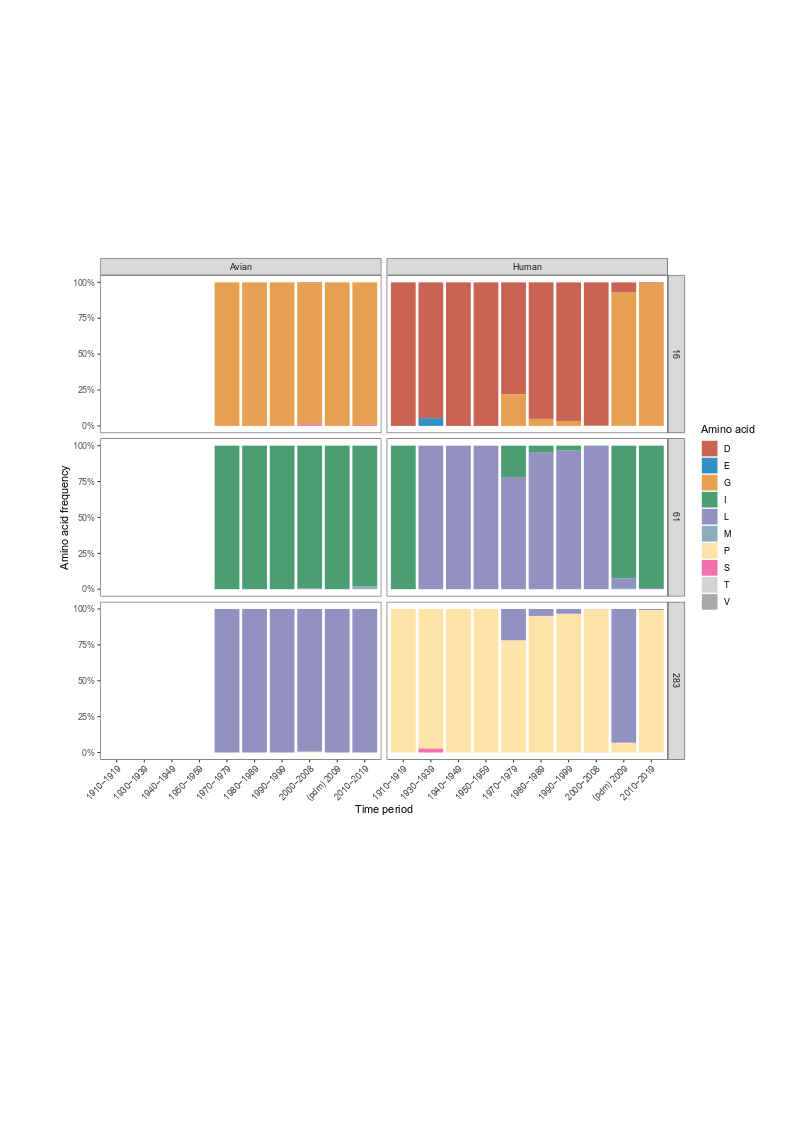

**Fig. S12. Amino acid frequency in the NP protein of human and avian H1N1 strains sorted by decade.** Displayed are the amino acid positions discussed in the main text (16, 61 and 283). Data source: Influenza Virus Database @NCBI, Filter: Complete NP sequences, H1N1, human versus avian. Downloaded on 2020-05-04**.** 1918 IAV sequences were not included.

| Primer | Sequence |
| --- | --- |
| BM PB2 fwd | AAAAGGTCTCAGGGAGCCGCCACCATGGAAAGAATAAAAGAACTAAGGG |
| MU PB2 fwd | AAAAGGTCTCAGGGAGCCGCCACCATGGAAAGAATGAAAGAACTAAGGG |
| PB2 rev | AAAAGGTCTCCTATTACTAATTGATGGCCATCCGAATTC |
| PB1 fwd | AAAAGCGATGAAGAATTCAAGGGAGCCGCCACCATGGATGTCAATCCGACTTTACTTTTC |
| PB1 rev | AAAAGCGATGAAGTATTCAATATTCTACTTTTGCCGTCTGAGCTC |
| PA fwd | AAAAGGTCTCAGGGAGCCGCCACCATGGAAGACTTTGTGCGACAATG |
| PA rev | AAAAGGTCTCCTATTCTATCTCAGTGCATGTGTGAGG |
| NP fwd | AAAAGGTCTCAGGGAGCCGCCACCATGGCGTCTCAAGGCACCAAACG |
| NP rev | AAAAGGTCTCCTATTTTAATTGTCGTACTCCTCTGCATTG |
| QC BM PB2-M631L fwd | CCACCAAAGCAAAGTAGATTGCAGTTCTCCTCTCTG |
| QC BM PB2-M631L rev | CAGAGAGGAGAACTGCAATCTACTTTGCTTTGGTGG |
| BM PB2 K627E fwd (for 631M) | CAGCCGCTCCACCAGAGCAAAGTAGAATGC |
| BM PB2 K627E rev (for 631M) | GCATTCTACTTTGCTCTGGTGGAGCGGCTG |
| BM PB2 K627E fwd (for 631L) | CAGCCGCTCCACCAGAGCAAAGTAGATTGC |
| BM PB2 K627E rev (for 631L) | GCAATCTACTTTGCTCTGGTGGAGCGGCTG |

#### Table S1. Cloning and mutagenesis primers for generation of polymerase complex expression constructs.

| Sample | Mapped reads | Endogenous human reads [%] | Average fragment length [nt] |
| --- | --- | --- | --- |
| MU162 | 1452871 | 6.53 | 62.7 |
| BE576 | 779471 | 4.28 | 59.8 |
| MU162/2 | 263258 | 1.01 | 118.3 |
| BE576/2 | 1795933 | 4.60 | 64.6 |
| BE572 | 123143 | 0.28 | 104.3 |

**Table S2. Mapping of the influenza-positive samples to a human transcriptome reference.** Shown are the results for independently generated libraries.

|  | **HSLC std** | **HSLC ext** |
| --- | --- | --- |
| **HA (w/ ambig) - only long** | 0 | .59 |
| **HA (w/ ambig)** | 0 | .99 |
| **HA (w/o ambig) - only long** | 0 | .62 |
| **HA (w/o ambig)** | 0 | 1 |
| **MP** | .75 | .78 |
| **NA** | .01 | .24 |
| **NP** | 1 | 1 |
| **NS** | .91 | .96 |
| **PA** | 1 | 1 |
| **PB1** | .44 | .68 |
| **PB2** | .92 | 1 |

Table S3. Support for human pandemic origin of the human seasonal influenza.

Numbers refer to the posterior probability that the respective segments from the human seasonal H1N1 lineages are direct descendants of the pandemic influenza.

| Gene | Strain | nt change | aa position | aa change |
| --- | --- | --- | --- | --- |
| HA | BE-576 | G356A | 119 | R119K |
|  | BE-576 | T1600C | 534 | Y534H |
|  | CU | G715A | 239 | D239N |
|  | CU | G472A | 158 | A158T |
| MP | BM | C10T | 4 | N |
|  | BE-572, MU | G543A | 181 | N |
|  | BE-572, MU | C693T | 231 | N |
|  | BM | C700A | 234 | L234I |
| MP2 | BM | T849G | 54 | L54R |
| NA | MU | A435G | 145 | N |
|  | BM | T462C | 154 | N |
|  | BE-572, MU | A588C | 196 | N |
|  | BM | C768A | 256 | F256L |
|  | MU | A774C | 258 | N |
|  | BM | C900T | 300 | N |
|  | MU | A1036G | 346 | I346V |
|  | MU | G1386A | 462 | N |
|  | BM | C1397G | 466 | T466S |
| NP | BE-572, BE-576 | A47G | 16 | D16G |
|  | BE-572, BE-576 | G165A | 55 | N |
|  | MU | A181T | 61 | I61L |
|  | BM | G504A | 168 | N |
|  | BE-572, BE-576 (only 2x coverage) | C848T | 283 | P283L |
|  | MU | A864G | 288 | N |
|  | BM | G987A | 329 | N |
|  | MU | C1221T | 407 | N |
|  | BM | T1488C | 496 | N |
| NS | MU | T64G | 22 | F22V |
|  | BM | G224A | 75 | G75E |
| PA | BE-572, BE-576 | T194C | 65 | S65F |
|  | BM | A453T* |  | N |
|  | BM | A456G* |  | N |
|  | BM | G473A* | 158 | R158K |
|  | BM | A495G* |  | N |
|  | BM | A498G* |  | N |
|  | BM | T501C* |  | N |
|  | BM, CU | A510G* | 170 | N |
|  | BM | G532A* |  | N |
|  | MU | C531T | 177 | N |
|  | MU, BE-572 | G1009T | 337 | A337S |
|  | MU | G1203A | 401 | N |
|  | CU | C1548T | 516 | N |
|  | BE-572 | T1557C | 519 | N |
|  | MU | C1709T | 570 | T570I |
|  | BM | C1710A | 570 |  |
| PB1 | BM | A161G | 54 | K54R |
|  | MU, BE-572, BE-576 | A1164G | 388 | N |
|  | MU | A2081G | 694 | N694S |
|  | MU | G2202A | 734 | N |
| PB2 | MU | A12G | 4 | I4M |
|  | MU, BE-572, BE-576 | C246T | 82 | N |
|  | MU, BE-572, BE-577 | G322A | 108 | A108T |
|  | MU, BE-572 | A552G | 184 | N |
|  | MU, BE-572 | A582G | 194 | N |
|  | MU, BE-572 | A639C | 213 | N |
|  | MU, BE-572 | G816A | 272 | N |
|  | BE-572 | T1013A | 338 | V338D |
|  | MU, BE-572 | A1038G | 346 | N |
|  | BE-572 | A1140G | 380 | N |
|  | MU | G1326A | 442 | N |
|  | MU | A1452T | 484 | N |
|  | MU | G1615A | 539 | V539I |
|  | MU | A1891T | 631 | M631L |

**Table S4. Nucleotide and amino acid changes in the pandemic strains analyzed.** Nucleotide (nt) and amino acid (aa) changes are reported with reference to the position in the coding sequence of the MU-162 strain (here referred to as MU). Each nt/aa change is reported as nt/aa observed in the other strains followed by the position and then the nt/aa observed in the strain that is different. N stands for no aa change conferred. * indicate nt differences identified in the potential recombinant fragment of the BM sequence.

| Library | Mapping reference | Median insert size (nt) | Maximum insert size (nt) |
| --- | --- | --- | --- |
| 162_1 | MU-162/1918 | 119.5 | 403 |
| 162_2 | MU-162/1918 | 118.5 | 200 |
| 162_3 | MU-162/1918 | 132.5 | 320 |
| 162_4 | MU-162/1918 | 133.5 | 236 |
| 162_5 | MU-162/1918 | 133 | 318 |
| 162_6 | MU-162/1918 | 112.5 | 184 |
| 162_7 | MU-162/1918 | 114.5 | 313 |
| 162_8 | MU-162/1918 | 115 | 337 |
| 162_9 | MU-162/1918 | 199 | 751 |
| 572_1 | MU-162/1918 | 166 | 532 |
| 572_1 | MU-162/1918 | 173 | 453 |
| 572_3 | MU-162/1918 | 201.5 | 381 |
| 572_4 | MU-162/1918 | 165 | 166 |
| 576_1 | MU-162/1918 | 133.5 | 306 |
| 576_2 | MU-162/1918 | 151 | 194 |
| 576_3 | MU-162/1918 | 133 | 281 |
| 576_5 | MU-162/1918 | 102 | 102 |
| 576_6 | MU-162/1918 | 113 | 113 |
| 576_9 | MU-162/1918 | 173.5 | 322 |

**Table S5. Median and maximum insert sizes for reads mapping to the 1918 influenza MU-162 genome.**
